# Algorithmic design of 3D wireframe RNA polyhedra

**DOI:** 10.1101/2022.04.27.489653

**Authors:** Antti Elonen, Ashwin K. Natarajan, Ibuki Kawamata, Lukas Oesinghaus, Abdulmelik Mohammed, Jani Seitsonen, Yuki Suzuki, Friedrich C. Simmel, Anton Kuzyk, Pekka Orponen

**Affiliations:** Department of Computer Science, Aalto University, 00076 Aalto, Finland; Department of Neuroscience and Biomedical Engineering, Aalto University, 00076 Aalto, Finland; Department of Robotics, Graduate School of Engineering, Tohoku University, Japan; Natural Science Division, Faculty of Core Research, Ochanomizu University, Japan; Physics Department E14, Technical University Munich, 85748 Garching, Germany; Department of Mathematics and Statistics, University of South Florida, Tampa, FL 33620, USA; Department of Applied Physics and Nanomicroscopy Center, Aalto University, 00076 Aalto, Finland; Frontier Research Institute for Interdisciplinary Sciences, Tohoku University, Japan; Division of Chemistry for Materials, Graduate School of Engineering, Mie University, Japan

**Author notes:** These authors contributed equally to this work (Antti Elonen, Ashwin K. Natarajan).

## Abstract

We address the problem of de novo design and synthesis of nucleic acid nanostructures, a challenge that has been considered in the area of DNA nanotechnology since the 1980s and more recently in the area of RNA nanotechnology. Towards this goal, we introduce a general algorithmic design process and software pipeline for rendering 3D wireframe polyhedral nanostructures in single-stranded RNA. To initiate the pipeline, the user creates a model of the desired polyhedron using standard 3D graphic design software. As its output, the pipeline produces an RNA nucleotide sequence whose corresponding RNA primary structure can be transcribed from a DNA template and folded in the laboratory. As case examples, we design and characterize experimentally three 3D RNA nanostructures: a tetrahedron, a bipyramid and a prism. The design software is openly available, and also provides an export of the targeted 3D structure into the oxRNA molecular dynamics simulator for easy simulation and visualization.

## Introduction

*Nucleic acid nanotechnology*, often more narrowly called *DNA nanotechnology*, uses nucleic acids as fabrication material for self-assembling nanoscale structures and devices (*1*). Major advances in this area, specifically in the self-assembly of structures, include *e*.*g*. the early multi-stranded DNA cube and truncated octahedron designs by Seeman *et al*. (*2, 3*), the mostly single-stranded DNA octahedron by Shih *et al*. (*4*), and the fundamental DNA origami technique by Rothemund (*5*) with its further applications to highly complex 2D (*6*–*8*) and 3D (*6, 7, 9*–*13*) designs.

While most current research in nucleic acid nanotechnology focuses on DNA-based nanostructures, there is also an emerging research tradition of using RNA as the fundamental substrate. One appeal of this alternative is that while the production of designed DNA nanostructures typically proceeds by a multi-stage laboratory protocol that involves synthesizing the requisite nucleic acid strands and hybridizing them together in a thermally controlled process, RNA nanostructures can, in principle, be produced in quantity by the natural process of polymerase transcription from a representative DNA template, isothermally at room temperature. The challenge, on the other hand, is that the self-assembly by folding of single-stranded RNA designs is not yet as robust and predictable as the self-assembly by hybridization of scaffold and staple DNA based on Rothemund’s origami technique (*5*).

The primary design methodology in this area of *RNA nanotechnology*, pioneered by Westhof, Guo, Jaeger *et al*. (*14, 15*) has been *RNA tectonics* (*16*–*18*), in which naturally occurring RNA motifs are assembled by connector motifs such as kissing hairpin loops (*17, 19*) or single-stranded sticky ends (*20*) into larger complexes. In a landmark article, Geary *et al*. (*21*) introduced the complementary approach of *RNA origami*, wherein a single long RNA strand is folded directly into a structure that conforms to a prescribed design. In this approach, kissing hairpin loops have a central role as connectors used to bring regions of the target structure together, but except for this use of the kissing loop motif, the method is *de novo*.^1^

The seminal article (*21*), together with its companion paper (*24*) on the detailed design principles, focused on the task of producing 2D “RNA origami tiles”, somewhat analogous to the DNA origami tiles introduced in (*5*). The basic methodology however carries within it a potential for similar extensions as those that followed the introduction of DNA origami in (*5*). A major step in this direction was taken by Li *et al*. (*25*), who presented designs and experimental characterizations of several 2D structures and a 3D tetrahedron. The versatility of the methodology was advanced by Liu *et al*. (*26*), who introduced, among other things, a novel “branching” kissing-loop connector motif that can be used to create trivalent branches in the structures, leading to a richer design space than was previously available. A further improvement was presented by Geary *et al*. (*27*) by enabling co-transcriptional folding of RNA origami of larger constructs compared to their earlier work (*21*).

In the present work, we contribute a solution to a broad family of further design challenges in RNA nanotechnology by providing a fully general design scheme and automated software pipeline for designing *arbitrary 3D RNA wireframe polyhedra*. Analogous schemes have been presented in recent years for 3D DNA wireframe structures (*12, 13, 28*), but because of the differences in the substrate, these multi-helix DNA origami based techniques do not carry over to RNA. To demonstrate our method, we display designs and experimental characterizations of three simple 3D wireframe structures: a tetrahedron (similar to (*25*)), a bipyramid and a prism.

## Results

### Structure Design

We describe our general design scheme using the simplest example, a tetrahedron (Fig. 1A-E). The starting point is a polyhedral model (Fig. 1A), whose wireframe representation we wish to render as a tertiary RNA structure (Fig. 1E).

**Fig. 1.**
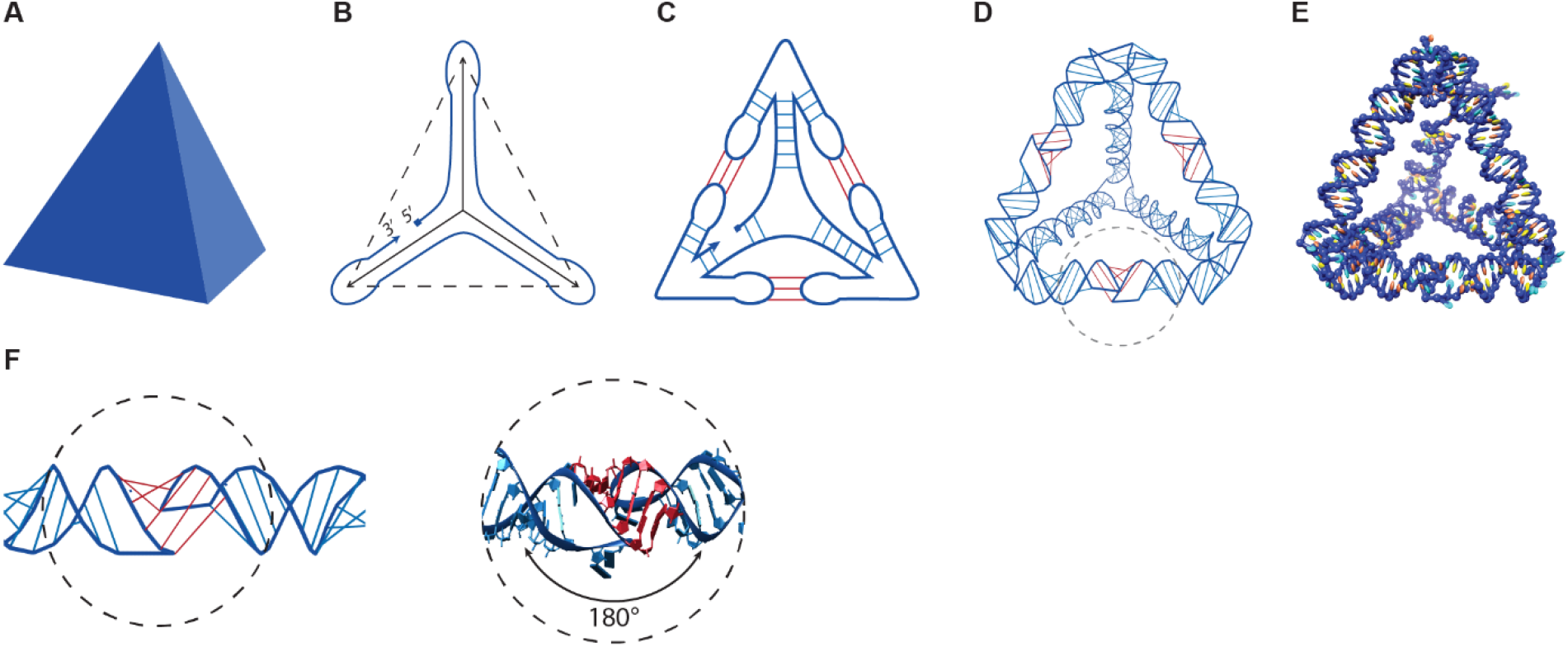
RNA polyhedron design scheme and HIV-DIS 180° kissing-loop. **(A)** Targeted polyhedral model; **(B)** Spanning tree and strand routing of the polyhedral mesh; **(C)** Routing-based stem and kissing-loop pairings; **(D)** Helix-level model and **(E)** Nucleotide-level model. **(F)** Schematic representation of the kissing loop and nucleotide-level model of the kissing-loop base pairing (Adapted with permission from (*21*)).

The first step is to create the RNA secondary structure of the targeted wireframe shape, deferring the precise nucleotide sequence design. We aim to render the edges of the wireframe mesh as RNA A-type helices, and towards that goal we wish to route the RNA strand around the edges of the mesh in such a way that every edge is covered twice by the routing, in antiparallel directions. With an appropriate nucleotide sequence design, the complementary strand segments will then hybridize together to create the duplex edges.

However, complete antiparallel double routings of polyhedral meshes that keep all vertices intact exist only under quite special conditions (*29*), and in particular for a tetrahedron such a routing is impossible. A way around this constraint is to first reduce the polyhedral mesh to one of its spanning trees,^2^ perform a strand routing on this tree, and then reintroduce the discarded edges using some connector motif.^3^ A schematic of the routing on a spanning tree of the tetrahedron mesh is presented in Fig. 1B, where the three solid lines indicate the chosen spanning tree edges, the three dashed lines the discarded non-spanning tree edges and the blue curve the routing of the strand around the spanning tree, with a nick between the 5’ and 3’ ends of the strand.

The connector we use to create the non-spanning tree edges is the 180° HIV-DIS kissing-loop motif successfully employed by Geary et al. in (*21*). This extrahelical pseudoknot pairing of two hairpin loops induces an almost perfect 180° alignment of the respective loop stems, and thus combines two separate “semi-helices” into an effectively contiguous helical structure (Fig. 1F). As outlined in Fig. 1C, we extrude the strand routing at every vertex of the mesh by a half-edge hairpin loop along each discarded non-spanning tree edge and design the base sequence at the loop terminus to pair with the corresponding half-edge that protrudes from the other end-vertex of the edge. The kissing-loop pairing of the matching termini will then reintroduce the non-spanning tree edges to the structure.

A small number of unpaired nucleotides are added to each vertex of the structure to provide flexibility and thus facilitate the folding of the 3D conformation. The exact number of these is determined by an optimization process that also tries to match the rotation phases of the helices incident to each vertex, so that the cross-vertex transition of the RNA strand from one helix to the next creates minimal strain to the conformation. (For more details of the design process, see Supplementary Text S1.)

Fig. 1D presents a helix-level model of the resulting RNA polyhedron, with the regular intrahelical nucleotide pairings marked in blue and the extrahelical kissing-loop pairings in red. A nontrivial aspect of the design process that is worth mentioning is the exclusion of strand routings that create mathematical knots and hence potential topological hindrances for successful folding.^4^ To ensure an unknotted routing, we imagine each vertex point as expanded into a small sphere, and choose for each incoming strand segment an outgoing segment in such a way that the connecting “virtual” routings on the surface of the sphere do not cross each other. This property can always be achieved by a judicious geodetic point-matching protocol and can be proved by a simple topological argument to guarantee that the resulting global routing is an unknot. (For details, see Supplementary Text S1. Because of the general way we treat the knotting problem, our design method actually covers not just polyhedra but even arbitrary straight-line 3D meshes.) We have developed a software tool *Sterna*^5^ (*31*) that automates the secondary-structure design process described above. This tool has been implemented as a Python add-on module to the open-source *Blender* 3D graphic design software suite (*32*). To initiate a design task, the user creates a model of the desired polyhedron using the tools in the *Blender* suite or imports an existing model from an external library. Pressing a “generate” button then performs the strand routing, creates the corresponding spatially embedded RNA A-helices, aligns their phases, and adds linker nucleotides at the vertices. The outcome can be viewed and edited on the Blender viewport screen or exported as a structured text file of type snac,^6^ which contains a representation of the resulting secondary structure in the standard “dots-and-brackets” notation, together with 3D coordinates of the nucleotides along the helices.

The snac file can then be carried to a further module *snacseq*, which complements it with kissing-loop base sequences optimized for binding strength and mutual orthogonality, and a full primary structure sequence designed with the help of the NUPACK package (*33, 34*). An additional module *snac2ox* can be used to transform this full snac file into input files for the *oxRNA* molecular dynamics simulation and visualization package (*35, 36*). (For further information, see Supplementary Note S1 and the software tutorial on the Sterna website (*31*).)

Fig. 1E illustrates the complete nucleotide-level model generated by the *Sterna* secondary structure design tool and the *snacseq* primary-structure creation module from the *Blender* design of a tetrahedron shown in Fig. 1A. The *snac2ox* module has been used to export the resulting design into *oxRNA*, which has then been used to relax the initial helix arrangement, and the output has been visualized using the *UCSF Chimera* package (*37*). Fig. 2 presents six further examples of *oxRNA* simulations of Sterna designs for RNA polyhedra – Bipyramid (Fig. 2A), Cube (Fig. 2B), Prism (Fig. 2C), Dodecahedron (Fig. 2D), Icosahedron (Fig. 2E) and Toroid (Fig. 3F) – illustrated in the same way.

**Fig. 2.**
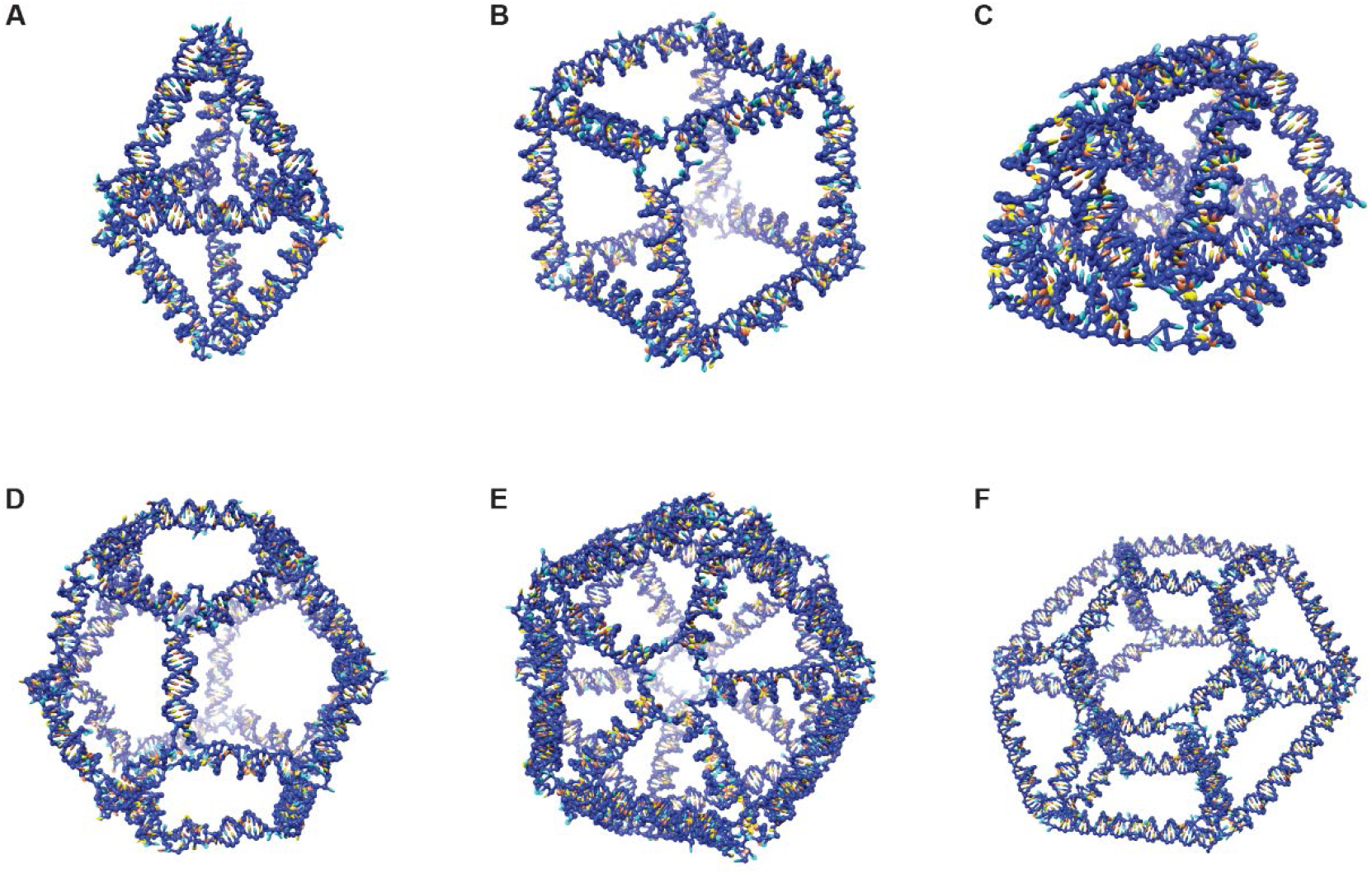
*oxRNA* models of Sterna designs for RNA polyhedra. **(A)** Bipyramid. **(B)** Cube. **(C)** Prism. **(D)** Dodecahedron. **(E)** Icosahedron. **(F)** Toroid.

**Fig. 3.**
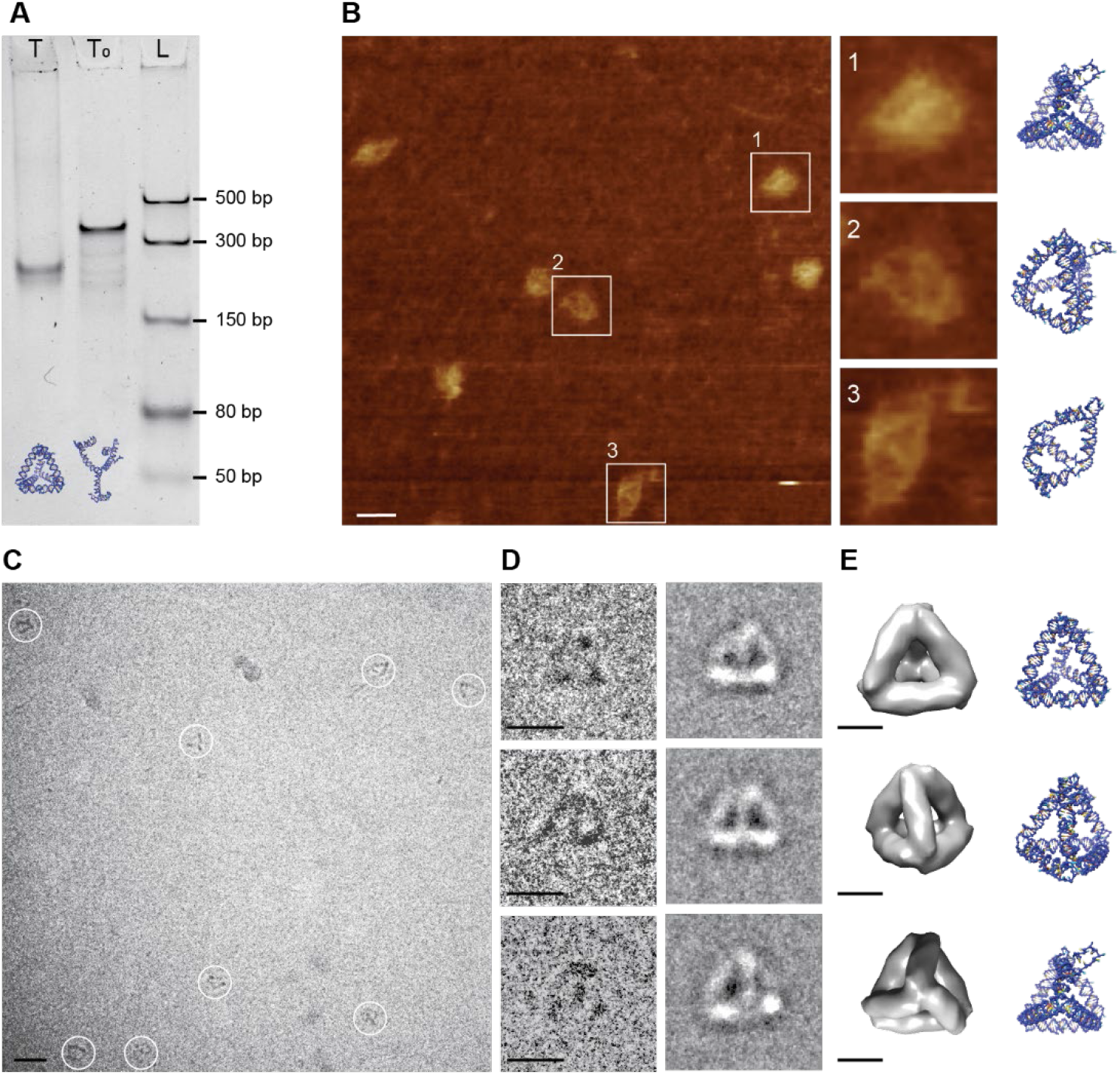
Characterization of Tetrahedron. **(A)** 5 % native PAGE analysis of tetrahedra T and T_0_ folded at 50 nM concentration. A significant difference in migration speed can be observed between T and T_0_, suggesting that T is folded completely into a compact tetrahedron by the interaction of kissing loops. **(B)** AFM micrograph of Tetrahedron T. The squares marking the particles and the enlarged images (right) are 30 × 30 nm in dimension. The enlarged images 1 and 2 on the right show the different orientations of tetrahedra on the surface while image 3 shows a tetrahedron with one kissing loop broken. Scale bar: 20 nm. **(C)** Cryo-EM micrograph of the Tetrahedron T sample. White circles indicate individual Tetrahedron particles; the circles have a diameter of 20 nm. The scale bar is 20 nm. **(D)** Individual particles picked from the micrographs (left) and their class averages (right). Scale bars: 10 nm. **(E)** Corresponding views of the Tetrahedron T structure reconstructed with 22 Å resolution from 1020 particles (left) and their *oxRNA* simulation models (right). Scale bars: 5 nm.

### Experimental Characterization

To validate our methodology, we synthesized three relatively small and distinctly characterizable 3D wireframe structures that were designed using the *Sterna* tool and the snacseq primary-structure generator: a Tetrahedron (Fig. 1E), a Bipyramid (Fig. 2A) and a Prism (Fig. 2C). (Note that the sides of the Prism structure have been triangulated to ensure structural rigidity.) In addition, the aforementioned structures with some or all kissing loops replaced with non-pairing sequences were used for comparing their folding efficiencies. The sequences and secondary structures of all the designs are presented in Supplementary Table S3 and Supplementary Fig. S7.

Tetrahedron (T) structures were chosen for the initial self-assembly for their simplicity compared to the other structures. The 453 nt long ssRNA for structure T was transcribed from the DNA template and purified using 8% denaturing PAGE. The purified ssRNA was folded by annealing in TE/2mM Mg^2+^/100 mM Na^+^ Buffer, resulting in formation of the hairpin duplexes and the three kissing loops. The native PAGE results show that a distinct construct presumed to be T folded at concentrations of up to 100 nM (Supplementary Fig. S8A). There was also an additional slow running band, which might be an aggregate of two or more T constructs. The tetrahedral structure T was compared with a deficient structure T_0_, where for each kissing loop one of the hairpin-loop sequences was replaced with a tetraloop sequence (*e*.*g*. GAAA) and the other with a poly-AU sequence (*e*.*g*. AUAUAU). Hence, the kissing loops do not close in T_0_ and the structures do not fold into tetrahedral conformations. 5% native PAGE analysis shows that T_0_ migrate distinctly slower than T, suggesting that the observed T are indeed completely folded Tetrahedra which are more compact than T_0_ (Fig. 3A).

The construct T was characterized further using atomic force microscopy (AFM) and cryo-electron microscopy (cryo-EM). AFM imaging performed on T indicates that the structures were completely folded and correspond to geometric tetrahedra in various projections (Fig. 3B). The enlarged AFM images 1 and 2 witness the tetrahedral symmetry of T, corroborating the PAGE results. The enlarged AFM image 3 shows a Tetrahedron structure with one broken kissing loop. We surmise that the AFM imaging itself causes the structures to open as the tip moves over the sample, which could explain the presence of partly unfolded structures observed in the image. Further confirmation of correct folding was provided by cryo-EM analysis. The initial cryo-EM imaging of T folded at 100 nM yielded an extremely low number of structures per frame (<1 particle per frame). Though 100 nM was a reasonable concentration for cryo-EM, the structures had an affinity towards the carbon in the grid, leaving very few structures in the hole. To increase the number of structures per frame, the sample was concentrated by spin filtering to ∼400 nM. The four-fold increase in concentration reduced the time and effort in data collection as the number of structures per frame increased significantly to about 3 particles per frame (Fig. 3C). In total, 1020 particles were picked, class averaged and reconstructed. The particles picked and their corresponding class averages show the different projections of construct T (Fig. 3D). The reconstruction revealed the tetrahedral structure of T with a resolution of 22 Å and the views corresponding to the class averages are presented in Fig. 3E. The reconstruction performed without imposing any symmetry (C1) also showed a tetrahedral structure, confirming that upon annealing the ssRNA of T would fold into a Tetrahedron via formation of the kissing loops (Supplementary Fig. S9A).

The Bipyramid (B) and Prism (P) structures were more complex, containing 5 and 7 kissing loops respectively, compared to 3 in structure T, which made characterization of these structures more challenging. The native PAGE analysis of B folded at different concentrations shows only little aggregation (Supplementary Fig. S8B). In addition to B with five kissing loops, we also folded control structures B_2_ and B_0_, in which three or all five of these kissing loops were replaced by tetraloop and poly-AU sequences. The construct B ran faster than B_2_ and B_0_, indicating the fully folded nature of B (Fig. 4A). The samples B_2_ and B_0_ migrated at the similar speed. This was somewhat unexpected since B_2_ still had two kissing loops intact, compared to none in B_0_. To understand this migration behavior, we calculated the radius of gyration (*k*) value for all the structures from the results of *oxRNA* simulation runs using Visual Molecular Dynamics (VMD). The mean values of *k* for structures T, T_0_, B, B_2_, B_0_, P and P_0_ were 5.33, 8.98, 6.29, 8.68, 9.27, 6.37 and 9.88 nm, respectively (Supplementary Fig. S10). We hypothesize that the close mean *k* values of B_2_ and B_0_ to be the reason for their observed migration speeds.

**Fig. 4.**
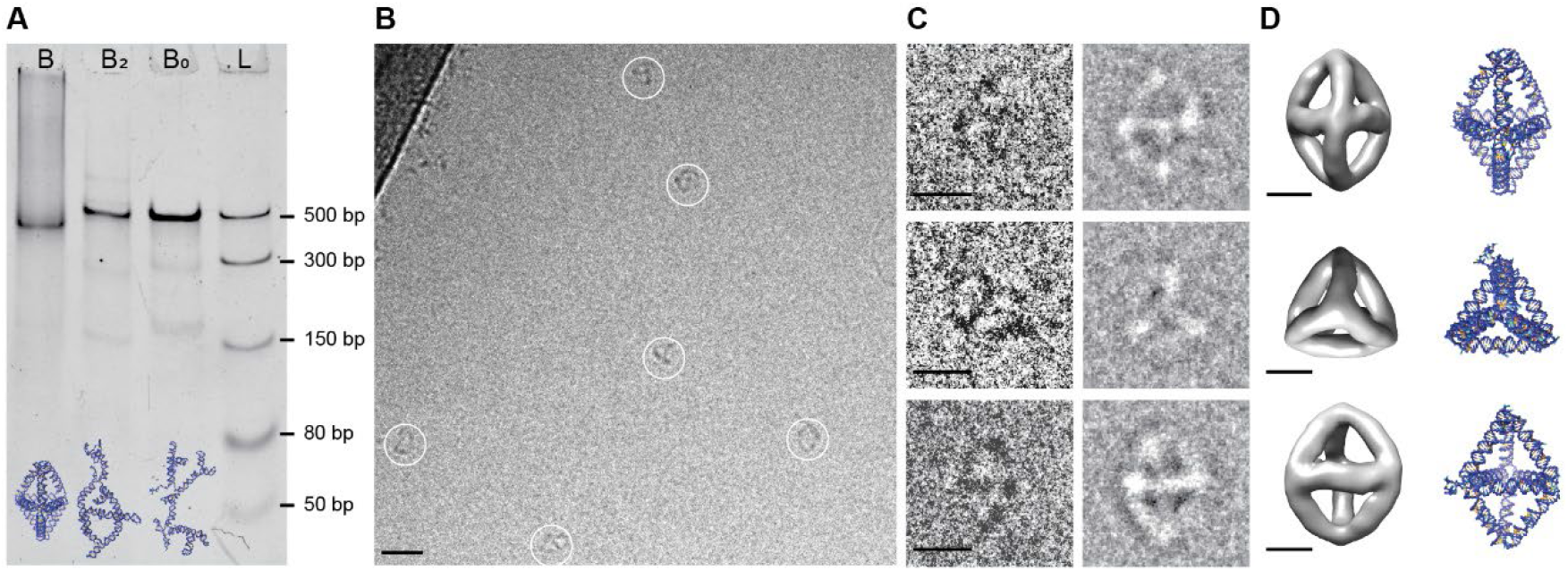
Characterization of Bipyramid. **(A)** 5% native PAGE analysis of Bipyramid B, partially deficient Bipyramid B_2_ and fully deficient Bipyramid B_0_. The faster running band at B indicates the folded Bipyramid. B_2_ and B_0_ run at similar speed. **(B)** Cryo-EM micrograph of the Bipyramid B sample. White circles indicate individual Bipyramid particles; each circle has a diameter of 20 nm. Scale bar: 20 nm. **(C)** Individual particles picked from the micrographs (left) and their class averages (right). Scale bars: 10 nm. **(D)** Different views of the Bipyramid B structure reconstructed from 955 particles (left) and their corresponding *oxRNA* simulation models (right). Scale bars: 5 nm.

The AFM imaging of the construct B did not yield any significant information due to the complexity of the structure. Hence, the construct was imaged under cryo-EM at ∼400 nM concentration (concentrated using spin filtering from samples folded at 100 nM). The number of structures per frame (∼2 particles/frame) was considerably lower than for the T structure (Fig. 4B). Individual particles were picked from the cryo-EM micrographs and class averaged. The particles picked and their class averages illustrate the different projections of the Bipyramid (Fig. 4C). The corresponding views of the construct B after reconstruction using 955 particles with a resolution of 25 Å are presented in Fig. 4D along with their *oxRNA* simulation models. The reconstruction of the Bipyramid without imposing any symmetry (C1) presented in Supplementary Fig. S9B also confirms the folding of the ssRNA template into a Bipyramid.

The Prism (P) structure contains seven kissing loops and folded without any aggregates at 10 nM concentration; however, at concentrations ≥ 20 nM, the sample started to aggregate and only a small amount of correctly folded structures migrated to form a faint band (Supplementary Fig. S8C). We compared the folding capability of P with that of the other two structures folded at 50 nM concentration by running the corresponding samples in parallel in a 5% native PAGE (Supplementary Fig. S8D). The constructs T and B folded without any aggregation while P had aggregates stuck in the gel pocket. Also, there was a second band close to the pocket which might be multimeric Prism structures formed by intermolecular kissing loop interactions. It could be observed that the migration speeds of T, B and P are determined by both the radius of gyration and the length of the ssRNA strand. Though B and P do not have a significant difference in radius of gyration (Supplementary Fig. S10), they migrate at different speeds because of their different strand lengths of 633 and 801 bases, respectively.

For comparison, a structure P_0_ with all the kissing loops replaced with tetraloops and poly-AU sequences was synthesized. The sample P_0_ ran as a single strong band, which was slower than sample P, indicating the fully folded nature of the P particles (Fig. 5A). We faced similar issues with AFM imaging of P as with B. The AFM could not resolve the structure and the images were noninformative. Our efforts to concentrate the Prism P sample with spin filtering also did not result in high yields, making cryo-EM imaging time consuming. A substantial portion of the structures found in the micrograph were either aggregated or had high affinity to the carbon in the grid. This resulted in fewer than 0.1 structures per frame on average. Some of the structures that were found intact and fully folded are presented in Fig. 5B, along with corresponding projections from the simulation model, suggesting the targeted prism structure was indeed realized.

**Fig. 5.**
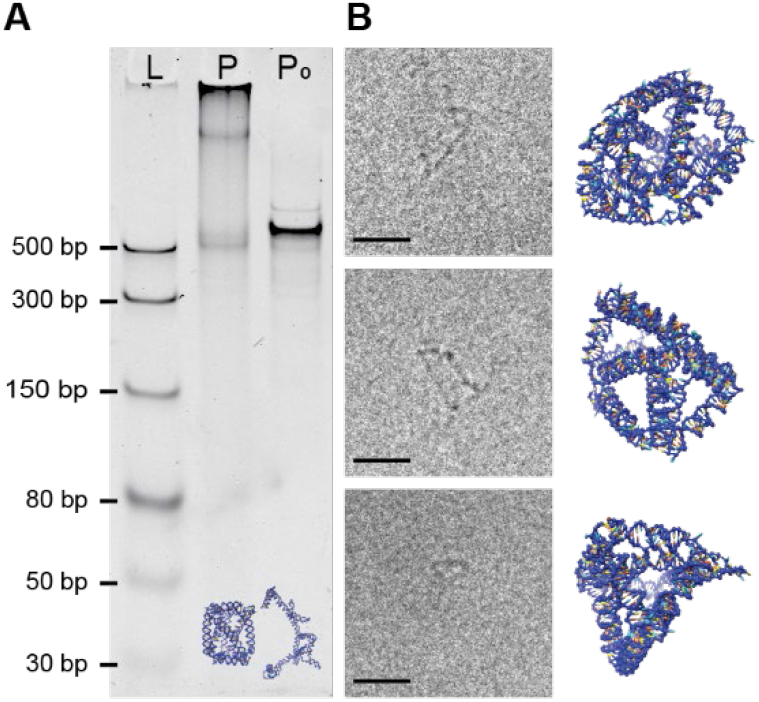
Characterization of Prism. **(A)** 5 % native PAGE analysis of Prism P and P_0_. The Prism P structures exhibited severe aggregation and a slow running band. The band running faster than P_0_ indicates the fully folded prism P. **(B)** Particles picked from Cryo-EM micrographs (left) and their corresponding *oxRNA* simulation views (right). Scale bars: 20 nm.

## Discussion

We have presented a general high-level and fully automated design scheme for rendering 3D wireframe polyhedra as native conformations of single-stranded RNA molecules. An open-source distribution of the design software is available at Sterna website (*31*). The method has been demonstrated by designing, synthesizing, and characterizing three small-to-moderate sized structures: a tetrahedron (455 bases), a bipyramid (663 bases) and a prism (801 bases). For the tetrahedron and the bipyramid the yield of the synthesis was high, and we were able to obtain excellent cryo-EM reconstructions. In the case of the prism a band of properly folded particles could be extracted, but the yield was too low to obtain decent quality reconstructions from the available number of cryo-EM grids. In further work we will investigate ways of counteracting this effect, taking into account the recent experimental advances of Li *et al*.(*25*), Liu *et al*. (*26*) and Geary *et al*. (*27*). We will also pursue the synthesis of bigger structures: in principle, our design approach sets no limits to the size or complexity of the target polyhedra, but in practice more attention will likely need to be devoted to RNA sequence design, in particular to the design of good kissing loop ensembles. This could be used to rationally counteract the observed aggregation tendencies and improve the robustness of the designs against possible defects in the nucleotide sequence. Utilizing RNA origami to functionalize components has been sparsely exploited. One remarkable example was introduced by Liu *et al*. (*38*), who utilized the kissing-loop interactions in RNA origami to functionalize target RNAs and assembled into closed homomeric nanoarchitectures for cryo-EM imaging of target RNA. Further developments in this direction should be explored to make RNA origami more robust and functional, similar to DNA origami.

## Materials and Methods

### Design of Sequences

Sequences for the RNA nanostructures excluding kissing loops were designed using the design function of NUPACK (*29, 30*). Six kissing loop sequences used for the design were taken from existing literature (*39*–*41*) and the rest were manually designed six base loops that are non-similar to others. (The snacseq tool provides a kissing-loop ensemble generator, but for the present experiments proven models from the literature, complemented with manual designs, were used to decrease the number of possible sources of difficulty.)

A restriction was imposed on the sequences to prevent any of the following patterns: AAAA, CCCC, GGGG, UUUU, KKKKKK, MMMMMM, RRRRRR, SSSSSS, WWWWWW, YYYYYY. In the corresponding complementary sequence, GT mismatches were introduced at 8 base pair intervals to avoid creating a self-complementary sequence. The mismatches were necessary to synthesize the DNA templates as dsDNA in the form of custom gBlock gene fragments from Integrated DNA Technologies (IDT, Inc.).

All the sequences begin with the cap sequence GAC followed by the T7 promoter sequence TAATACGACTCACTATAG. The cap and promoter sequences are also used as a sequence for forward primer during PCR. The designs also have a 15nt tail sequence in addition to the sequence required for the nanostructure. Different 15nt tail sequences were used for Tetrahedron, Bipyramid, and Prism, with the intention of using the 15nt sequence as a reverse primer.

However, to optimize the melting temperature (*T*_*m*_) of forward and reverse primers, slightly longer reverse primer sequences were used. See Supplementary Note S2 for the detailed script to design the sequence of DNA template using NUPACK.

The DNA templates for all RNA nanostructures were ordered from IDT, Inc. as custom gBlock gene fragments. The primer sequences for PCR amplification were ordered from IDT, Inc. All the DNA sequences and the primers used can be found in Supplementary Table S3.

### DNA amplification and RNA synthesis

The gBlocks from IDT were resuspended in nuclease free water and the final concentration was 10 µg/µl. The DNA templates were amplified over 20 cycles using Phusion High-Fidelity DNA Polymerase (New England Biolabs, Inc.). The final concentration of components in a 100 µl PCR reaction was 1ng/µl of DNA template, 0.5 µM of forward and reverse primers and 1x Phusion High-Fidelity PCR Master Mix with HF Buffer. The reaction mixture was denatured initially at 98 °C for 30 s, followed by cycles of 8 s of 98 °C, 15 s of 58 °C and 30 s of 72 °C. The final elongation step was held for 7 min at 72 °C. The PCR product was purified using Monarch PCR & DNA Cleanup Kit (New England Biolabs, Inc.). The samples were run on a 1.5% Agarose (Merck) gel with SyBr Safe (ThermoFisher Scientific) in 0.5x TBE buffer at room temperature for 60 minutes at 120 V. The gels were imaged using Bio-Rad Gel Doc XR system. The purified PCR DNA templates (∼20 ng/µl) were transcribed in a reaction mixture containing 4 mM of each rNTP, 12.5 mM of MgCl_2_, 1x RNA polymerase buffer and ∼1 U/µl of T7 RNA polymerase (New England Biolabs, Inc.). The samples were incubated at 37 °C for 6 hours, and an additional hour after the addition of 2 U/ 50 µl of DNase (New England Biolabs, Inc.). The RNA samples were purified using an 8% Acrylamide:Bis-Acrylamide (29:1) containing 8 M Urea and 1x TBE. The samples were preheated at 95° C for 5 minutes and loaded on the gel. The samples were run at 58 °C for 100 min at 100 V. The gel was post stained with SyBr Green (ThermoFisher Scientific) for 15 min and imaged using Bio-Rad Gel Doc XR system. The bands of interest were cut from the denaturing gels and purified using ZR small-RNA PAGE Recovery Kit (Zymo Research). The purified RNA samples were stored in nuclease free water at -20 °C.

### Assembly of RNA nanostructures and characterization

The RNA samples were thermally annealed in 0.5x TE buffer with 1 mM MgCl_2_ and 100 mM NaCl. 100 nM of ssRNA in TE/Mg^2+^/Na^+^ buffer was rapidly folded by heating to 80 °C for 5 min followed by cooling to 20 °C at 0.1 °C/s. The samples were run in a 5% native PAGE gel in an ice bath along with low range ssRNA ladder (New England Biolabs, Inc.). The gels were prepared using Acrylamide: Bis-Acrylamide (29:1) containing 1 mM MgCl_2_ and 100 mM NaCl. The gels were post stained with SyBr Green (ThermoFisher Scientific) for 15 min and imaged using Bio-Rad Gel Doc XR system.

### AFM imaging

Atomic force microscopy (AFM) imaging was performed using a tip scan high-speed AFM (BIXAM, Olympus, Tokyo, Japan) that was improved based on a previously developed prototype AFM (*42, 43*). A freshly cleaved mica surface was pre-treated with 0.05% 3-aminopropyltriethoxy silane (APTES) (*44*). A drop (2 µL) of the sample (about 1 nM) in the TAE-Mg buffer (40 mM Tris-acetate, 1 mM EDTA, 2mM Mg2Cl, pH 8.3) was deposited onto the APTES-treated mica surface and incubated for 3 min. The surface was subsequently rinsed with 10 µL of the TAE-Mg buffer. Small cantilevers (9 µm long, 2 µm wide, and 100 nm thick; USC-F0.8-k0.1-T12, NanoWorld, Switzerland) having a spring constant of ∼0.1 N/m and a resonant frequency of ∼300–600 kHz in water were used to scan the sample surface. The 320 × 240-pixel images were collected at a scan rate of 0.2 frames per second (fps). The imaged sequences were analyzed using an AFM scanning software (Olympus) and ImageJ (http://imagej.nih.gov/ij/) software.

### Cryo-EM imaging and single particle reconstruction

For cryo-EM, 100 nM of RNA polyhedra samples folded in TE/Mg^2+^/Na^+^ buffer were concentrated four times using 3 kDa MWCO Amicon Ultra centrifugal filters (Merck). 5 µl of the concentrated sample was applied on 300 mesh Cu grids coated with lacey carbon (Agar Scientific). The grids were blotted for 3s (70% relative humidity) by Leica EM GP2 plunge freezer followed by immediate vitrification using liquid ethane (−170 °C). Vitrified samples were cryo-transferred to the microscope and imaged using a JEOL JEM-3200FSC TEM while maintaining specimen temperature of 86 K. EMAN2 (*45*) was used for single particle reconstruction of the RNA nanostructures. The reconstruction of tetrahedron was performed with 1020 particles that were used for reference free class averaging. The initial models were generated by imposing cyclic C1 (no symmetry) or tetrahedral symmetry (TET) and refined with C1, C3 and TET symmetry. For Bipyramid reconstruction, 955 particles were picked for class averaging, initial reconstruction and refinement. The reconstructed models were visualized using UCSF Chimera.

## General

IK thanks Kaori Tanabe for the help with lab work in Tohoku University. We acknowledge the provision of facilities and technical support by Aalto University at OtaNano - Nanomicroscopy Center (Aalto-NMC) and the computational resources provided by the Aalto Science-IT project.

## Funding

Academy of Finland grant 311639 (AE, AM and PO)

Academy of Finland grant 308992 (AKN and AK)

Japan Society for the Promotion of Science (JSPS) Early-Career Scientists 18K18144 (IK)

Fund for the Promotion of Joint International Research (B) 19KK0261 (IK)

Young researcher dispatch program (School of Engineering, Tohoku University) (IK)

NSF-DMS grant numbers 1800443/1764366 (AM)

Nokia Foundation-2017 (AM)

JSPS Grant-in-Aid for Scientific Research KAKENHI-grant numbers 18K19831 and 19H04201 (YS)

European Research Council grant agreement no. 694410, project AEDNA(LO and FCS)

## Author contributions

Conceptualization: PO, IK and AM initiated the study.

Methodology: PO introduced the strand routing scheme, with the use of kissing loops as connector motifs suggested by FCS. IK and LO did the initial strand designs. The Sterna design tool was developed and implemented by AE, with contributions from AM and PO. Later strand designs were done using the Sterna tool, with detailed adjustments by IK. Investigation: Experimental designs were discussed together, with laboratory work divided between TU Munich (Initial RNA transcription and folding experiments; LO & FCS), Aalto University (Optimization of RNA transcription, folding and sample preparation by AKN & AN) and Tohoku University (RNA transcription, folding, and sample preparation for AFM; IK & YS).

Computations for gyration radii of the RNA structures were performed by IK.

Visualization: Cryo-EM imaging (AKN, AK & JS) and AFM imaging (AKN, IK & YS)

Numerical simulations and helix-level visualizations were done by AE.

Supervision: PO, AK, FCS.

Writing—original draft: The main manuscript was primarily written by AKN and PO, with significant contributions from IK, AK and JS. The description of the Sterna design tool in Supplement 1 was primarily written by AE, with contributions from PO.

Writing—review & editing: PO, AKN, AK, FCS.

## Data and materials availability

All data needed to evaluate the conclusions in the paper are present in the paper and/or the Supplementary Materials.

## Footnotes

1 Besides these two general approaches, one may mention also the work of Afonin et al. (*22*), where RNA cubes are constructed from a small number of intertwined short RNA strands, and that of Han et al. (*23*), where 2D RNA tiles of various shapes are created by locking antiparallel overlays of partially-complemented regions together with cohesive parallel crossover connections.

2 In graph theory, a *spanning tree* of a graph *G* is a cycle-free set of edges that connects all the vertices of G (*30*, Section 1.5).

3 This design technique for single-stranded constructs goes back at least to Shih and co-worker’s mostly single-stranded DNA octahedron (4) and has resurfaced many times, although the connection to spanning trees was apparently made explicit first by Veneziano et al. (*13*). For an overview, see (*28*).

4 When discussing knottedness, we consider the strand to be a closed loop where the 3’-to-5’ nick is sealed. In practice the RNA strand of course folds from an open conformation, so knotting is not an absolute obstacle, but nevertheless might lead to kinetic traps in the folding process.

5 Short for “*S*panning *T*ree *E*ngineered *RNA* design”.

6 Short for “*S*imple *N*ucleic *A*cid *C*ode”.

