## Supplementary Materials for "Algorithmic design of 3D wireframe RNA polyhedra"

**This PDF file includes:**

Supplementary Text
Figs. S1 to S10
Tables S1 to S3
References (46 to 53)

**Supplementary Text**

S1 Algorithmic design methods

In this Supplementary Note we describe in detail our algorithmic methods for designing an RNA primary structure that folds into a target polyhedral model. We first discuss the task of strand routing, then helix generation, and finally sequence generation.

The methods apply in principle to any straight-line 3D mesh (that is, a connected graph linearly embedded in three-dimensional Euclidean space), but for simplicity we frame the discussion in terms of polyhedra, or more precisely polyhedral *beam-frameworks*, i.e., the edge systems of polyhedra considered as rigid beams. The structures of arguably greatest interest are the convex polyhedra, but also nonconvex ones or ones of higher genus (e.g., toroids) can be considered.

For practical purposes it is often desirable that the polyhedron is structurally *rigid*. It is well known that a convex polyhedron (or rather its beam-framework) is rigid if and only if the polyhedron is fully triangulated, i.e. all its faces are triangles (*46*). Also for nonconvex polyhedra triangulation seems to be in practice helpful for improving rigidity, but theoretically the issue is highly nontrivial.

S1.1 Strand routing

As discussed in the main paper, our strand routing for a given target polyhedron *P* is initiated by choosing a spanning tree *T* for *P*[1] and tracing a path around *T* in such a way that every edge in the tree is covered twice, in antiparallel directions, by the routing. This can be achieved by a simple "twice-around-the-tree" walk, widely used in graph algorithms (*47*) and often even in DNA nanostructure design either explicitly or implicitly (*28*).

Because the basic twice-around-the-tree walk on *T* only covers the edges in *P* included in the spanning tree, we extend the walk from each vertex $v$ of *P* also halfway down each non-spanning tree edge incident to $v$, creating a hairpin loop structure (Fig. S1). The terminal loops of the two antipodal half-edge hairpins along each edge $\{u, v\}$ are paired by a kissing-loop design.

Given a desired scaling of the polyhedron *P*, this routing scheme induces an RNA secondary structure design that has a native conformation in the shape of *P*, when augmented with a matching primary sequence where appropriate care has been taken to address the risk of nonspecific bindings and other sequence-design issues.

It is however not guaranteed that there is always a smooth folding pathway for the eventual strand to fold into the desired conformation. Firstly, the choice of the spanning tree *T* may affect the pathway, with some tree structures resulting in more reliable folding than others.

Secondly, and even more importantly, it is possible that the chosen twice-around-the-tree walk on *T* creates a topological *knot* (*48*, *49*), when one considers the walk as a closed circuit with the 3'-5' nick in the strand sealed. Fig. S2 illustrates how the basic trefoil knot can be created even in the routing of the simple three-pointed star spanning tree of a polyhedron in Fig S1 if one is not careful in choosing the traversal order of edges incident to each vertex.

One might think that this problem can be avoided by always traversing the edges in the order induced by a planar embedding of *T* as in Fig. S1. This is however not true, because the polyhedron *P* is in fact a spatial structure and embedding its spanning tree *T* in a plane may involve arbitrary choices on the relative ordering of the edges. Thus, a routing that is nicely

[1] More precisely, the beam-framework graph of *P*. However, for simplicity we will mostly not make this distinction.

uncrossing in a chosen planar embedding of T may nevertheless create a knot when lifted back to the spatially embedded $P$.[2]
Because an RNA strand is not a closed loop but a linear thread, knottedness of the routing is not a mathematically insurmountable topological obstacle to folding. Since it nevertheless may present a serious kinetic trap, we prefer to avoid it in our detailed routing method. This is done by controlling the traversal order of edges at each vertex by a *geodetic ordering* scheme, described in more detail below.
At a general level, creating a twice-around-the-tree routing proceeds as a simple depth-first traversal of the given spanning tree $T$ (*47*):

1. Choose some edge $e_0 = \{u_0, v_0\}$ of $T$ as the root edge and initialize the routing path as $\pi = [u_0, v_0]$. Perform traversal step 2 with initial edge $e = e_0$, oriented as $(u_0, v_0)$.
2. Let the initial edge be $e = (u, v)$. Sort all edges of $T$ incident to $v$ in geodetic order (to be defined) starting from $e$. Let the sorted order of the edges be $e, e_1, \dots, e_n$. For each edge $e_i = \{v, v_i\}$, in this order, add vertex $v_i$ to the end of path $\pi$ and perform step 2 recursively with $e_i = (v, v_i)$ as the initial (oriented) edge. Finally add vertex $u$ to the end of path $\pi$.
3. Repeat traversal step 2 with $e = e_0$ oriented as $(v_0, u_0)$ as the initial edge.
4. Return path $\pi$.

As we shall see, to achieve an unknotted strand routing it suffices that the strand segments constituting the edges of our polyhedron $P$ do not form a "tangle" in the local geometric neighborhood of any vertex. To discuss this issue further we need to note, firstly, that the spanning tree $T$ is in fact embedded in space as indicated by the polyhedron $P$, and secondly that to control the local tangling, we also need to consider the half-edges that will be used to create the non-spanning tree edges of $P$. Thus, the twice-around-the-tree algorithm above actually needs to be run on the "extended" spanning tree $\hat{T}$ that contains these half-edges in addition to those in T, as presented in Fig S1.
To address the goal of keeping the local strand routing untangled, we imagine a small sphere $S_v$ drawn around each vertex $v$ of $P$ and think of each edge $e$ incident to $v$ as landing on $S_v$ at a point $x_e$. (Actually, since each edge is made up of two strand segments, we would have two landing points $x'_e$ and $x''_e$. But we can think of these as being arbitrarily close to each other and also arbitrarily rotatable, so that we can ignore this detail.) Now if the traversal order of the edges at $v$ is $e_0, e_1, \dots, e_n$, we can consider their strand connection pattern as following the piecewise-linear, closed curve $x_{e_0} - x_{e_1} - \dots - x_{e_n} - x_{e_0}$ on the surface of $S_v$. If this curve does not cross itself, then by the Jordan-Schönflies theorem it bounds a (topological) disk and hence is an *unknot* (*49*).
Thus, we can ensure a good arrangement of the strand routing at a vertex $v$ by any edge-traversal scheme which guarantees that the local edge-to-edge connectivity curve on $S_v$ is non-crossing. A simple way of achieving this is to first order the landing points $x_e$ on the surface of $S_v$ by their geodetic coordinates: fix some polar axis direction for $S_v$ (e.g., according to the first edge entering $v$), assign to each landing point $x_e$ a longitude and a latitude, and sort the points by longitude; or in the case that two points have exactly the same longitude then by latitude. Then traverse the edges in this order. (For convenience, one can also zig-zag the traversal order by alternating increasing and decreasing latitude orders between longitude groups.) This traversal scheme is illustrated in Fig S3.
Let us now establish that having locally non-crossing edge traversal orders at each vertex guarantees that also the global strand routing is an unknot. Towards this goal thicken each edge $e$ to a narrow cylinder and think of the two strand segments forming the edge as traversing on the

[2] This is not a problem if $P$ is convex, or more generally if the 3D mesh we are considering can be embedded on an orientable surface. Then the edges incident to each vertex do have a natural clockwise (say) ordering induced by the orientation on the ambient surface.

opposite sides of this cylinder. (Now the edges of course meet the surfaces of the vertex spheres at two points rather than one, but as discussed, we can always adjust the locations of these landing points so that they do not introduce crossings to otherwise non-crossing local traversal curves.)
By this scheme we achieve a globally non-crossing, closed strand-routing curve on the surface of the "thickened" extended spanning tree $\hat{T}$ composed of the vertex spheres and their connecting edge cylinders. But the surface of a thickened tree is topologically a sphere, so by another application of the Jordan-Schönflies theorem, this curve is also an unknot.
A minor point that still maybe requires clarification is that since the edges in the eventual RNA structure are construed as RNA double helices, it is not immediately clear that the constituent strands can be abstracted as traversing on the "opposite sides" of a helix cylinder. Could the helical spiraling be an additional source of knottedness? A mathematical argument to show that the answer is "no" is that despite the spiraling, the resulting strand-routing curve is still non-crossing, and so the Jordan-Schönflies theorem applies. A more intuitive explanation might be that the helical spirals can be unwound to straight lines by counter-rotating the helix cylinders and readjusting the strand-sphere landing points. (Or alternatively by counter-rotating whole subtrees of the thickened tree $\hat{T}$, starting from its leaves.)

<u>S1.2 Helix generation</u>
The helix generation step determines the positions of nucleotides along the previously determined strand-routing path. First, we choose a scale factor, which defines the exact lengths of the polyhedral edges in nanometers, and thus determines how many bases are placed on each edge. This factor must be large enough to allow for kissing loops to form. A kissing loop requires at least 13 bases, and so the path segment corresponding to such a loop must have room for at least 13 bases. Once a valid scaling factor has been chosen, we can use the default P-stick model parameters (*21*) to place the beads on the path.
Next, we determine how close to a vertex the incident helices can be placed without creating overlaps between them. This can be thought of as placing a sphere of radius R on the vertex so that no helix segments outside of the sphere overlap with each other. We determine the value of R for each vertex by Equation 1 below, where $\alpha$ is the most acute angle between edges incident to that vertex and $r$ is the radius of a helix in the P-stick model:

$$R = \frac{r}{\tan \alpha} \qquad (1)$$

As the helices defined by the P-stick model can rotate around their axes, it is not quite enough to know only the routing path and the scale factor, but we must also decide on good rotation angles for the helices. Poor rotational alignment might result in long gaps between consecutive helices or even in strand tangles. This problem is illustrated in Fig. S4.
To determine the alignment, double helices can be modeled as cylinders with four anchor bases linking them to other helices. Each cylinder can then be assigned a four-dimensional vector $l$, whose elements are the lengths of the links incident to it. Now, the problem of aligning the helices becomes one of minimizing some appropriate norm of each of these vectors by rotating the cylinders. The norm $||l||_2$ would minimize the sum of the link lengths, whereas the norm $||l||_\infty$ would minimize the length of the longest link. (Our Sterna tool uses norm $||l||_5$ by default, but this is an input parameter that can be adjusted.) To do this minimization, we use a "spring relaxation" algorithm, which tries to iteratively optimize the rotation phase of each cylinder while the other cylinders stay fixed.
First, we calculate the exact positions of the anchors for each cylinder by using the P-stick model parameters listed in Table S1(*21*). The first anchor will be at one end of the cylinder with an angle of 0. The second anchor will be $(N-1) \cdot D$ nanometers from the first anchor at angle $(N-1) \cdot$

$T$, where $N$ is the number of base pairs in the helix, $D$ is the helix rise per base pair and $T$ is the helix twist per base pair. The third and fourth anchors are the same as the first and second, but they are rotated by $A$ degrees and shifted by $I$ nanometers, where $A$ is the axis angle across the minor groove and $I$ is the base inclination relative to the helix axis. The chosen scale factor is very important here, since the twist $T$ for A-form helices is 32.73 degrees, which means that the angle repeats every 11 steps. Therefore, the anchors at both ends of a cylinder with $11n$, $n \in \mathbb{Z}$, bases will have the same angle. Such cylinders are easy to orient, and a shape consisting of only them is likely to have a very good optimal solution.
The second step of the spring relaxation is to calculate the torque exerted upon each anchor by the other cylinders. We can do this one anchor at a time by projecting the anchor of the other cylinder onto the normal plane of the current cylinder and by determining the angle $\gamma$ between the current position of the anchor and the projection. The formula for this is shown in Equation 2, where $x$ is the position vector of the current anchor, $d$ the position vector of the other anchor relative to the current anchor and $e$ is the unit axis vector of the cylinder. The exact torque is proportional to this angle. We can next add the signed angles of each anchor together to get the total torque exerted upon the cylinder. We then rotate the cylinder accordingly and proceed to the next cylinder.

$$\gamma \;= \arccos\left(\frac{x^T(d-(d^Te)e)}{||x||\,||d-(d^Te)e||}\right) \tag{2}$$

Once all the cylinders are optimally aligned, we can proceed by placing the nucleotides on their surfaces as described by the P-stick model. Since the distance between nucleotides serving as anchors for two consecutive helices will usually be longer than that between two bases along a regular helix, we must add extra linker bases between the anchor bases. This is not a problem if the number of linkers stays small, e.g., fewer than four, since we would normally add at least one linker between helices anyway in order to allow some flexibility at the vertex junctions. If there are too many linkers, however, the structure may start to lose its rigidity.
If a cylinder corresponds to a kissing-loop edge, we generate the nucleotides only halfway along the length of the edge and then turn back. The second half is formed when we revisit the edge from the opposite direction later. Because we want the kissing loops to serve the same function as double helices, they must form a 180-degree angle in relation to each other. This can be achieved by placing two unpaired bases before the loop and one after it as depicted in Fig. S5 (*21*). This also keeps the kissing loops consistent with the P-stick model. Since we use 180-degree kissing loops, we can treat also the kissing loop edges as regular cylinders in the spring relaxation phase. The outcomes of this design process are the (targeted) secondary and tertiary structures of the folded RNA strand. The final step is to design the primary structure sequence.

S1.3 Sequence design

We use the NUPACK (*33*) software package as our main sequence design tool. However, since NUPACK does not support the design of kissing loops, we generate them manually and use NUPACK only for designing the “plain” part of the structure that does not contain pseudoknots. NUPACK uses a constrained multistate test tube design algorithm, which optimizes the primary structure based on a nearest-neighbor model. The primary structure of the designed RNA in IUPAC nucleic acid notation is generated from its corresponding secondary and tertiary structure in three steps: designing the non-pseudoknotted plain part, choosing kissing loops and generating the eventual sequence.

S1.3.1 Designing the plain part

We define the *plain* part of an RNA structure as the bases that constitute the nested base pairs, the unpaired bases and the linkers, according to the given secondary structure. Bases that belong to any possible pseudoknots do not belong to this part.

Since RNA strands are transcribed from a single-stranded (ssDNA) template, there are limitations on sequence design that are induced by potential obstacles to the template synthesis, such as ssDNA secondary structure. As is conventional, we avoid the following sequences (in nucleic acid notation): AAAA, CCCC, GGGG, UUUU, KKKKKK, MMMMMM, RRRRRR, SSSSSS, WWWWWW and YYYYYY (*24*). Some further restrictions were imposed by other constraints in the supplier's synthesis process.

In the transcription process (T)hymine and (G)uanine in DNA are transcribed to (U)racil and (G)uanine in RNA, respectively. U's and G's form strong base pairs in RNA whereas T's and G's repel each other. By exploiting this difference in base pairing characteristics, we design sequences that form secondary structures in RNA but not in the complementary template ssDNA. Similarly as in (*24*), we replace every X:th base pair in the plain part with either U-G or G-U, thereby reducing secondary structure formation in ssDNA. Since the optimal value for X is hard to predict, we simply replace in this way every sixth base pair, rounding up when necessary. Thus, we prevent unwanted secondary structures in the template DNA while still retaining freedom in the sequence design. In nucleic acid notation, K's correspond to G's and U's, so we replace every sixth base pair in the plain part simply with K's.

In addition to base pairs, the plain part consists of unpaired bases and linker bases between consecutive helices, added to provide flexibility at the vertex junctions (Fig. S6). We carefully design the linkers to avoid base pairing with adjacent bases in the plain part. Considering that G's and C's could potentially create base pairs, and U's also form strong bonds with G's, we typically use A's in the linker region wherever possible. However, in structures with more than three consecutive linkers, using only A's would violate one of the restrictions described earlier. Hence, a combination of A's and U's (W's in nucleic acid notation) is used in these places.

The forward and reverse *primers* needed for the *polymerase chain reaction* are designed at 15 bases each with the following sequences: "GACUAAUACGACUCACUAUAGGG" and the corresponding secondary structure "....................(((" for the forward primer at the 5' end, and "NNNNNNNNNNNNNNN" and corresponding secondary structure "..............." for the reverse primer at the 3' end.

S1.3.2 Kissing loops

As described earlier, we use manually designed kissing loop sequences that are added to the plain part as fixed bases. Kissing loops are marked as unpaired bases in the secondary structure to avoid base pairing with the plain part. The kissing loop sequences used in the present designs were all six bases long, with the loops flanked by the sequences UGAA on the 5'-side and ACG on the 3'-side. The A's were added as padding to be certain that the loops are oriented at a 180 degree angle when they create a kissing-loop complex by base pairing (*21*).

The nearest neighbor model allows for the calculation of the energy between any two paired strands. If kissing-loop base pairs are assumed to be energetically similar to regular base pairs, strong kissing-loop pairs that are *orthogonal* to each other can be identified using nearest neighbor energetics. Since two strands pairing up will always have some binding energy in the nearest neighbor model, the conventional definition of orthogonality needs to be adapted. In the present work, two kissing-loop pairs were considered to be orthogonal to each other, if the mismatch pairing energies of their strands were below a certain threshold relative to the intended pairings.

Kissing loop sequences that pair only with their intended complementary loops can be designed either from scratch or by utilizing existing sequences from literature. The RNA polyhedra designs in the present work utilize kissing loops from the literature (*24*, *50*). However, since there are only a handful of reported kissing loop pairs, this would be a limitation for larger structures that require multiple such pairs.

Hence, we also considered the problem of designing new kissing loop pairs. For a loop of six bases, there are $4^6 = 4096$ possible sequences. A kissing loop pair consists of two (semi-) complementary loops, hence there are $4096^2 = 16777216$ possible pairs in total. We calculated the energies for all these possible pairs using the nearest neighbor energetics in NUPACK and sorted them in decreasing order according to their binding energies. The top ten kissing loop pairs are listed in Table S2.

This family of all possible six-base kissing loop pairs could also be considered as a weighted complete graph, where the loop sequences are the nodes and the edge weights the binding energies of each kissing loop pair. The problem of finding $n$ good kissing loop pairs now becomes one of finding a perfect matching of $2n$ nodes where all the edges forming the matching have weight below a certain $E_{pair}$, and all the other edges between the matched nodes have weights above a certain $E_{mis}$. Finally, minimize for $E_{pair}$ and maximize for $E_{mis}$. Although finding efficient ways of doing this remains an interesting challenge, the following naive approach may be helpful:

1. Sort all kissing-loop pairs in increasing order according to their binding energies (i.e. strongest bindings first). Mark all their constituent strands as "unselected". Choose a mispairing energy threshold $E_{mis}$.
2. Start processing the list from the top down until you have marked $2n$ strands as "selected".
3. Calculate the binding energies between both strands of the currently considered pair and all strands previously marked as "selected". If both values are higher than $E_{mis}$, mark both strands as "selected".
4. Return the $n$ strongest kissing loop pairs constituted from the $2n$ "chosen" strands. If the end of the list was reached before $2n$ strands were chosen, the algorithm has failed for the given parameter settings.

### S2 Sequence generation

Once all nucleotides are adequately restricted, the final RNA nucleotide sequence is generated using NUPACK. NUPACK identifies one or more strands where the number of mispaired bases is below a user defined threshold. If the structure is too constrained, NUPACK will fail since no sequences reach the necessary threshold.

In our experiments, after the primary structures were generated, we further validated them by computer simulations. We used IPknot to calculate the minimum nearest-neighbor energy model conformations for the structures, and checked that the expected secondary structures were produced (*51*) (Fig. S7). Other tools, such as Kinefold, SimRNA and oxRNA (*36*, *52*, *53*) were used to simulate the actual molecules in coarse-grained physical representations to predict the folding pathways and the final tertiary structures. Since this latter approach works only for very small structures within reasonable timeframes due to the extremely large search spaces, it is not suitable for strands longer than a few hundred nucleotides. We used the validated primary structures for DNA strand synthesis and further transcribed them to RNA. The DNA sequences for the RNA polyhedral structures and their corresponding primer sequences are listed in Table S3.

#### S2.1 DNA Templates and Secondary Structure Diagrams

NUPACK scripts used to design the DNA templates for the experiments:

##### S2.1.1 Tetrahedron

**---from here---**

material = RNA

temperature[°C] = 26.0
trials = 10
sodium[M] = 1.0
dangles = some
allow mismatch = true
structure RNA_seq =
...................(((((((((((((((..(((((((((((.........)))))))))))..(((((((((((.........)))))))))))..)))))))))))))))(((
((((((((((((..((((((((((((((((((((((((((((((((..(((((((((((.........)))))))))))..(((((((((((.........)))))))))))..))))
))))))))))))))))))))))))))))..((((((((((((((((((((((((((((((((..(((((((((((.........)))))))))))..(((((((((((.........))
)))))))))..))))))))))))))))))))))))))))))))..)))))))))))))))...............
domain a =
GACUAAUACGACUCACUAUAGGGNNGNNNNNUNNNNNAANNNNGNNNNNUGAAG
GAGGCACGNNNNNUNNNNAANNNNGNNNNNUGAAGCGAGCACGNNNNNUNNNNA
ANNNNNGNNNNNUNNCCCNNNNUNNNNNUNNNNNAANNNNNUNNNNNNGNNNNN
NGNNNNNNUNNNNNNAANNNNUNNNNNUGAAGCUCGCACGNNNNNGNNNNAANN
NNGNNNNNUGAAGUGGACACGNNNNNUNNNNAANNNNNNGNNNNNNUNNNNNNU
NNNNNNGNNNNNAANNNNNUNNNNNNGNNNNNNUNNNNNNGNNNNNNAANNNNU
NNNNNUGAAGUCCACACGNNNNNGNNNNAANNNNUNNNNNUGAAGCCUCCACGNN
NNNGNNNNAANNNNNNUNNNNNNGNNNNNNUNNNNNNGNNNNNAANNNNNGNNN
NNGNNNNNNNNNNNNNNNNNNN
RNA_seq.seq = a
prevent = AAAA, CCCC, GGGG, UUUU, KKKKKK, MMMMMM, RRRRRR, SSSSSS,
WWWWWW, YYYYYY
**---to here---**

S2.1.2 Bipyramid
**---from here---**
material = RNA
temperature[C] = 26.0
trials = 10
sodium[M] = 1.0
dangles = some
allow mismatch = true
structure RNA_seq =
...................(((((((((((.........)))))))))))..((((((((((((((((((((((((((((((((..((((((((((((((((((((((((((((((((..(
((((((((((.........)))))))))))..(((((((((((.........)))))))))))..))))))))))))))))))))))))))))))))..(((((((((((.........)
))))))))))..(((((((((((.........)))))))))))..))))))))))))))))))))))))))))))))..((((((((((((((((((((((((((((((((..((((
(((((((.........)))))))))))..(((((((((((.........)))))))))))..))))))))))))))))))))))))))))))))..((((((((((((((((((((((
((((((((((..(((((((((((.........)))))))))))..(((((((((((.........)))))))))))...(((((((((((.........))))))))))).)))))))))
)))))))))))))))))))))))...............
domain a =
GACUAAUACGACUCACUAUAGGGUNNNNNNUGAAAGGCUCACGNNNNNNGCCCAA
NNNUNNNNNGNNNNNUNNNNNGNNNNNUNNNNNAANNNUNNNNNGNNNNNGNNN
NNUNNNNNUNNNNNAANNNUNNNNNNUGAAGAGCCUACGNNNNNNGNNNAANNN
GNNNNNNUGAAGCGAGCACGNNNNNNUNNNAANNNNNGNNNNNGNNNNNUNNNNN
NUNNNNNGNNNAANNNUNNNNNNUGAAGCCUGCACGNNNNNNGNNNAANNNUNN
NNNNUGAAGUGGACACGNNNNNNGNNNAANNNNNGNNNNNUNNNNNGNNNNNUN
NNNNGNNNAANNNUNNNNNGNNNNNGNNNNNUNNNNNUNNNNNAANNGNNNNN
NUGAAGUCCACACGNNNNNNUNNNAANNGNNNNNNUGAAGGAGGCACGNNNNN
NUNNNAANNNNNGNNNNNGNNNNNUNNNNNUNNNNNGNNNAANNNUNNNNNUNN

NNNGNNNNNGNNNNNUNNNNNAANNNUNNNNNNNUGAAGCCUCCACGNNNNNNGN
NNAANNNGNNNNNNUGAAGCAGGCACGNNNNNNUNNNAAANNNGNNNNNNUGAA
GCUCGCACGNNNNNNUNNNANNNNNGNNNNNUNNNNNUNNNNNGNNNNNGNNNN
NNNNNNNNNNNNNNN
RNA_seq.seq = a
prevent = AAAA, CCCC, GGGG, UUUU, KKKKKK, MMMMMM, RRRRRR, SSSSSS,
WWWWWW, YYYYYY
**---to here---**

<u>S2.1.3 Prism</u>
**---from here---**
material = RNA
temperature[C] = 26.0
trials = 10
sodium[M] = 1.0
dangles = some
allow mismatch = true
structure RNA_seq =
...................((((((.........))))))..((((((.........)))))).((((((((((((((((.........)))))))))))))))).(((((((((((((((((((
(((((((((((...((((((((((((((((((((((.((((((((((((((((((.........))))))))))))))))))..(((((((((((.........)))))))))))...(((
(((.........))))))..)))))))))))))))))))))).((((((((((((((((((((..((((((((.........)))))))).(((((((((((((((((.........)))))))))
))))))).(((((((((((((((((((((((((((((((...((((((((((((((((((((((.((((((((((((((((((.........))))))))))))))))))..(((((((
(((((.........)))))))))))...(((((((.........))))))).)))))))))))))))))))))))..(((((((.........))))))).((((((((((((((((.........)
))))))))))))))).))))))))))))))))))))))))))))))))...))))))))))))))))))))).((((((((((((((((((.........))))))))))))))))))
..)))))))))))))))))))))))))))))))...............
domain a =
GACUAAUACGACUCACUAUAGGGNUGAAGUCACCACGNCCCAANNNNUGAAGCGU
UCACGNNNNANNNNGNNNNNNGNNNUGAAAGGCUCACGNNNUNNNNNNUNNNNA
NNNUNNNNNGNNNNNUNNNNNGNNNNNUNNNNNAAANNNNGNNNNNNUNNNNNN
UNNNANNNUNNNNNGNNNNNNUGAAGAGCCUACGNNNNNNUNNNNNGNNNAANN
NUNNNNNNUGAAGCGAGCACGNNNNNNGNNNAAANNNNUGAAGCCUGCACGNNNN
AANNNGNNNNNNGNNNNNNUNNNNANNNNUNNNNNNUNNNNNNUNNNAANNNNN
UGAAGCAGGCACGNNNNNANNNNUNNNNNNNGNNUGAAGUGGACACGNNUNNNN
NNNGNNNNANNNUNNNNNGNNNNNGNNNNNUNNNNNUNNNNNAAANNNNUNNNN
NNGNNNNNNUNNNANNNUNNNNNNUNNNNNNUGAAGUCCACACGNNNNNNGNNNN
NGNNNAANNNUNNNNNNUGAAGCUCGCACGNNNNNNGNNNAAANNNNNUGAAGA
ACGCACGNNNNNANNNGNNNNNNUNNNNNNGNNNNAANNNNNUGAAGGUGACAC
GNNNNNANNNNGNNNNNNNGNNUGAAGGAGGCACGNNUNNNNNNNUNNNNANNN
NNGNNNNNGNNNNNUNNNNNNUNNNNNGNNNAAANNNGNNNNNNGNNNNNNGNNN
NANNNUNNNNNNUNNNNNNUGAAGCCUCCACGNNNNNNGNNNNNGNNNAANNNNN
GNNNNNUNNNNNGNNNNNUNNNNNGNNNNNNNNNNNNNNNNN
RNA_seq.seq = a
prevent = AAAA, CCCC, GGGG, UUUU, KKKKKK, MMMMMM, RRRRRR, SSSSSS,
WWWWWW, YYYYYY
**---to here---**

<u>S3 Radius of Gyration</u>
The tcl script to compute radii of gyration from oxRNA simulations is shown below.

puts $frames

```
set radiusG [open "radiusG.txt" w]
for {set i 0} {$i <$frames} {incr i} {
 animate goto $i
 set sel [atomselect top "all"]
 set com [veczero]
 foreach coord [$sel get {x y z}] {
  set com [vecadd $com $coord]
 }
 set com [vecscale [expr 1.0/[$sel num]] $com]
 set sum1 0
 foreach coord [$sel get {x y z}] {
  set sum1 [expr $sum1 + [veclength2 [vecsub $coord $com]]]
 }
 set rg [expr sqrt($sum1 / ([$sel num] + 0.0))]
 puts $radiusG "$i $rg"
}
flush $radiusG
close $radiusG
```

The computed values for radii of gyration are plotted in Fig. S10.

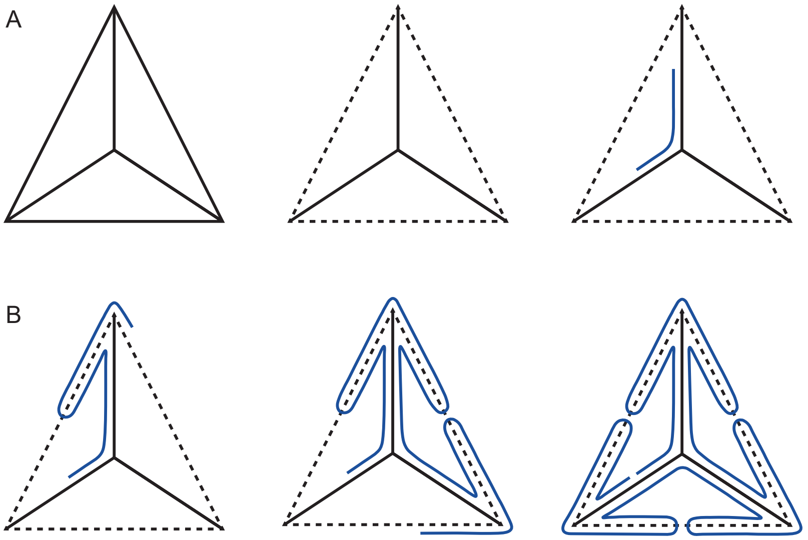


**Fig. S1: The twice-around-the-tree strand routing algorithm.** A: Beam-framework graph *G* of a tetrahedron; a spanning tree *T* of *G*; initial segment of strand route along *T*. B: Extruding the route along a non-tree edge; intermediate stage of route; complete route.

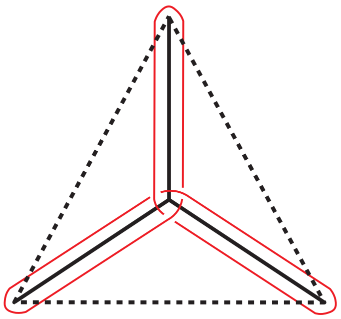

**Fig. S2: A trefoil-knot routing of a simple spanning tree.**

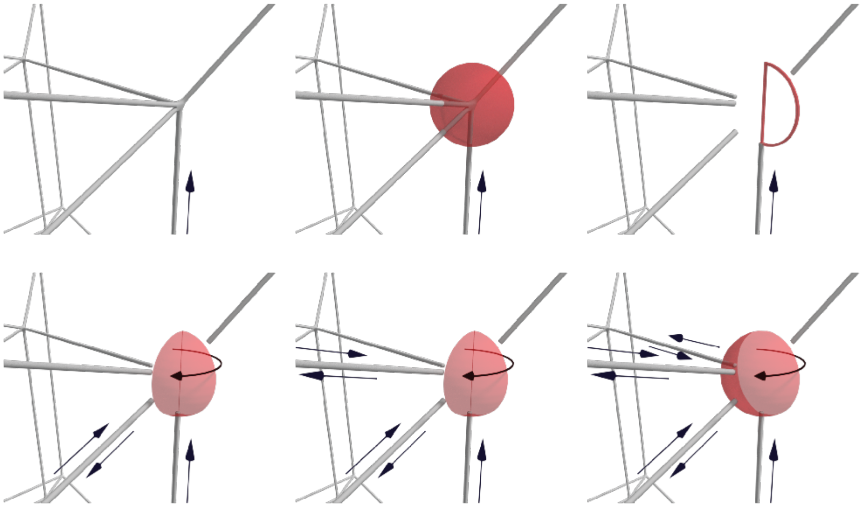

**Fig. S3: The geodetic vertex routing algorithm.**

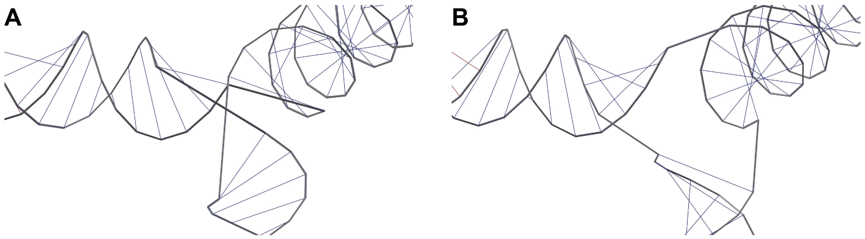


**Fig. S4: An example of a malformed vertex junction and a well-formed junction.** In the left image all three helices are poorly oriented, resulting in a tangle at the junction. In the second image all helices are well oriented.

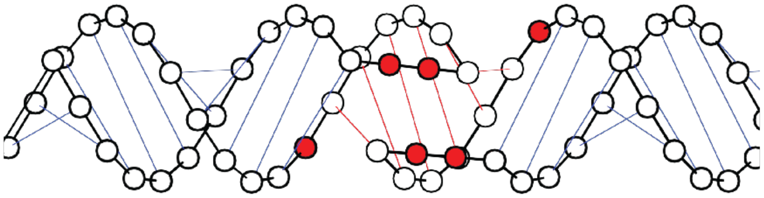

**Fig. S5: A kissing-loop pair.** The unpaired bases around the pseudoknot are marked red.

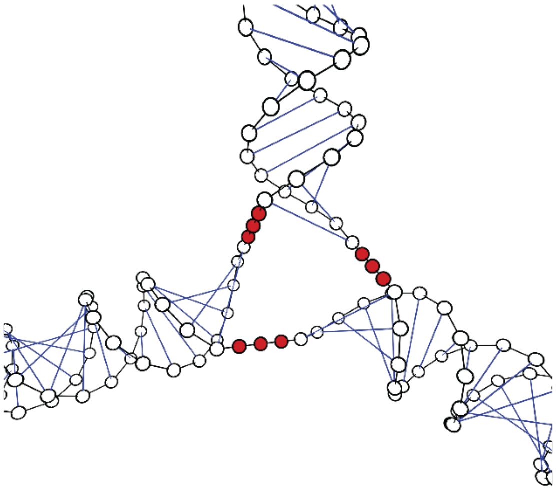

**Fig S6: A junction between three helices.** Linker bases are portrayed as red beads.

**A**

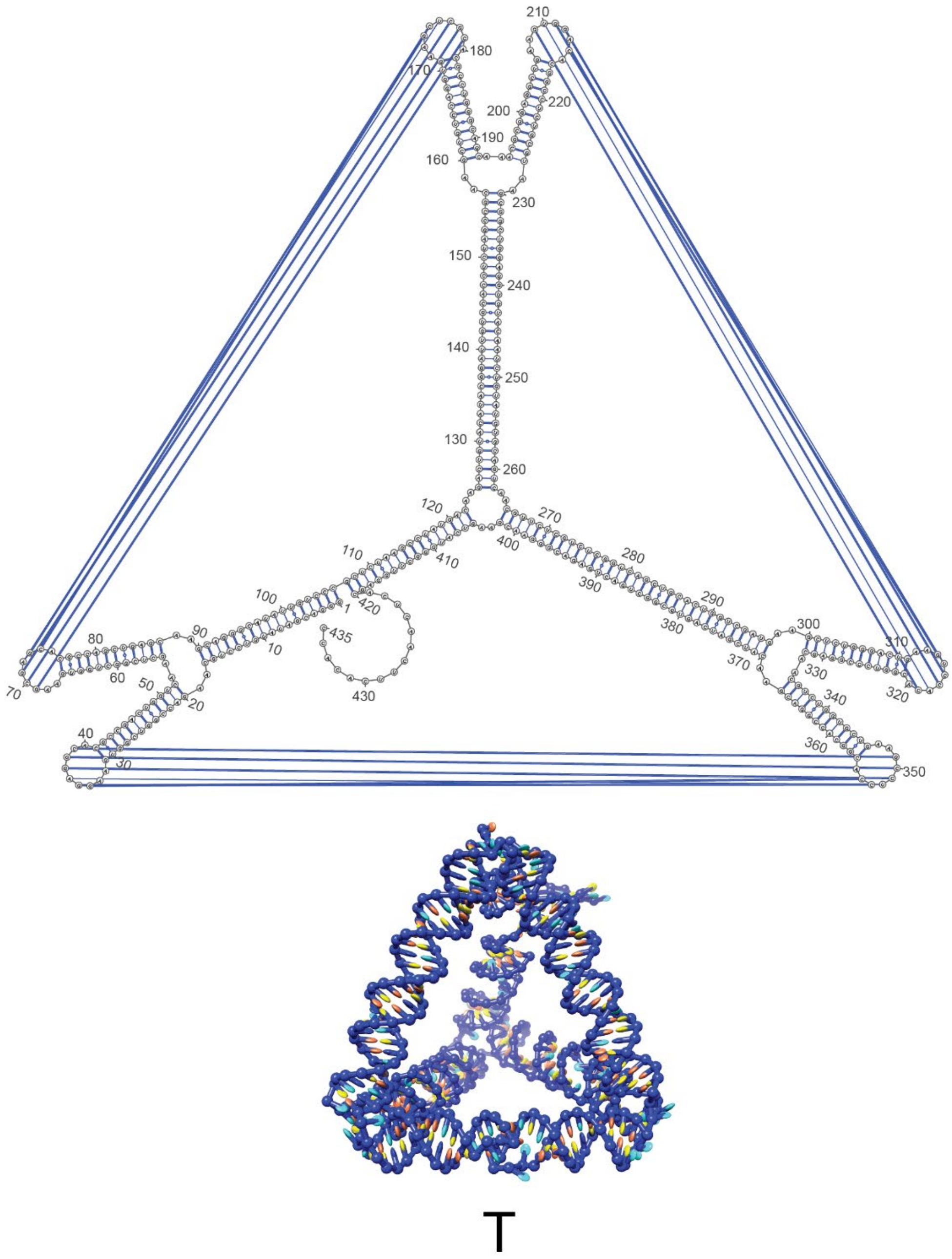

**B**

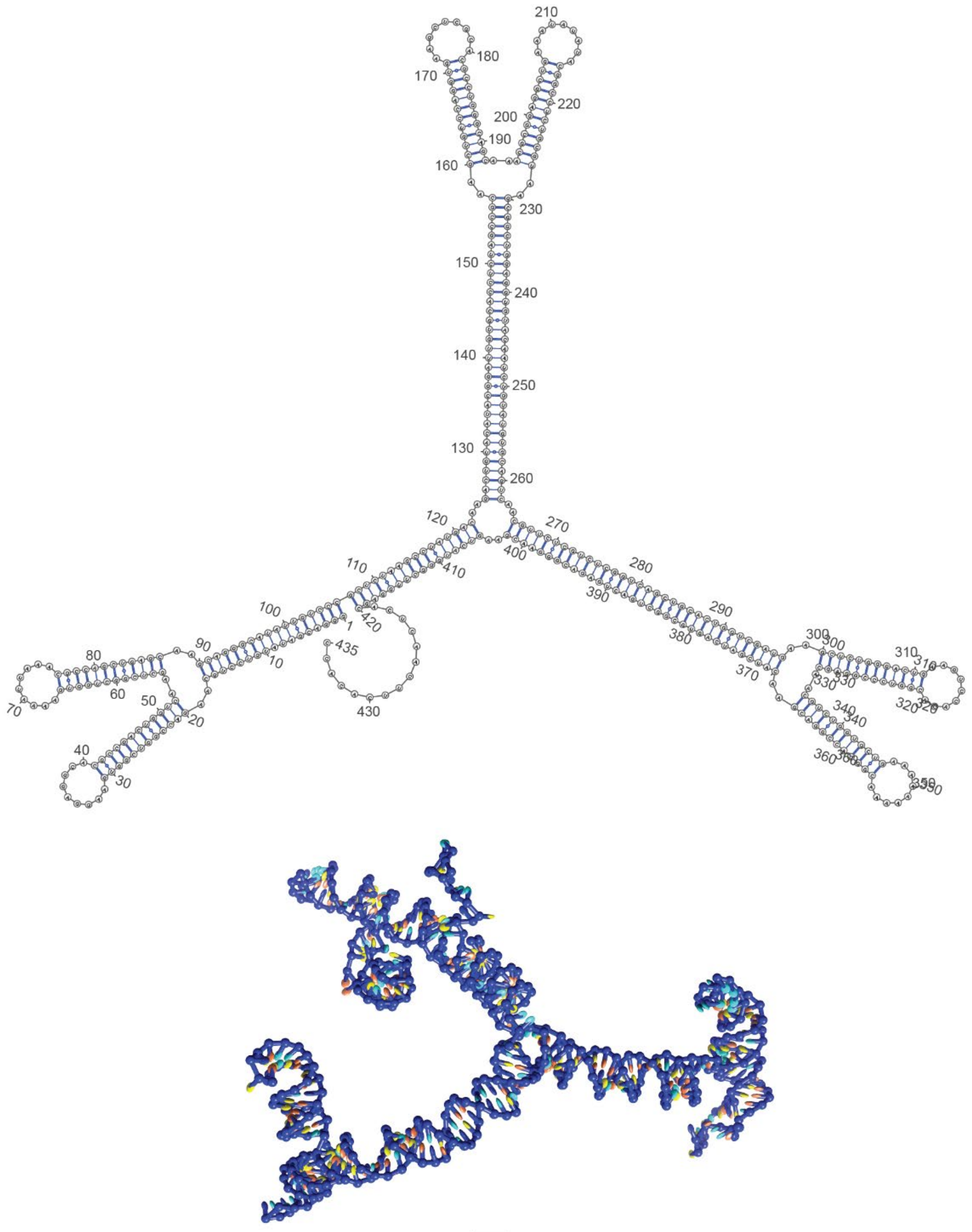

210
180
170
220
200
190
160
230
150
240
140
250
130
260
120
270
110
400
410
280
390
100
420
1
435
290
380
80
90
10
300
310
370
60
70
50
20
430
330
340
320
40
30
360
350


T₀

c

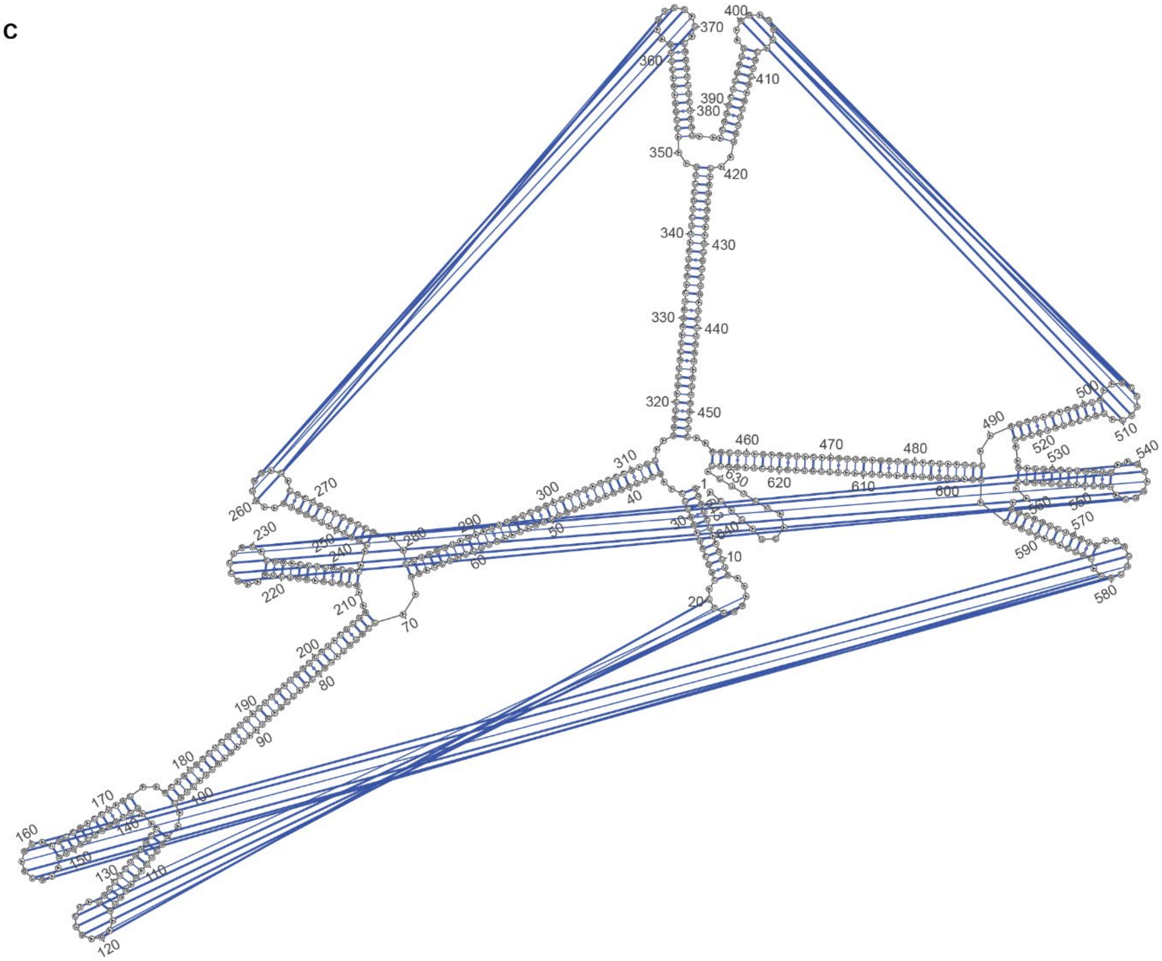
370
400
360
410
390
380
350
420
340
430
330
440
320
450
460
470
480
490
500
510
540
520
530
310
300
290
270
260
230
220
210
200
190
180
170
160
130
120
110
100
90
80
70
60
50
40
30
20
10
1
630
620
610
600
570
580
590

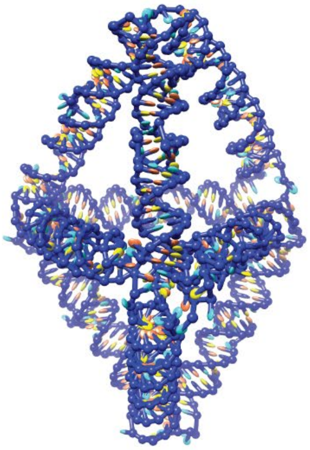

B

**D**

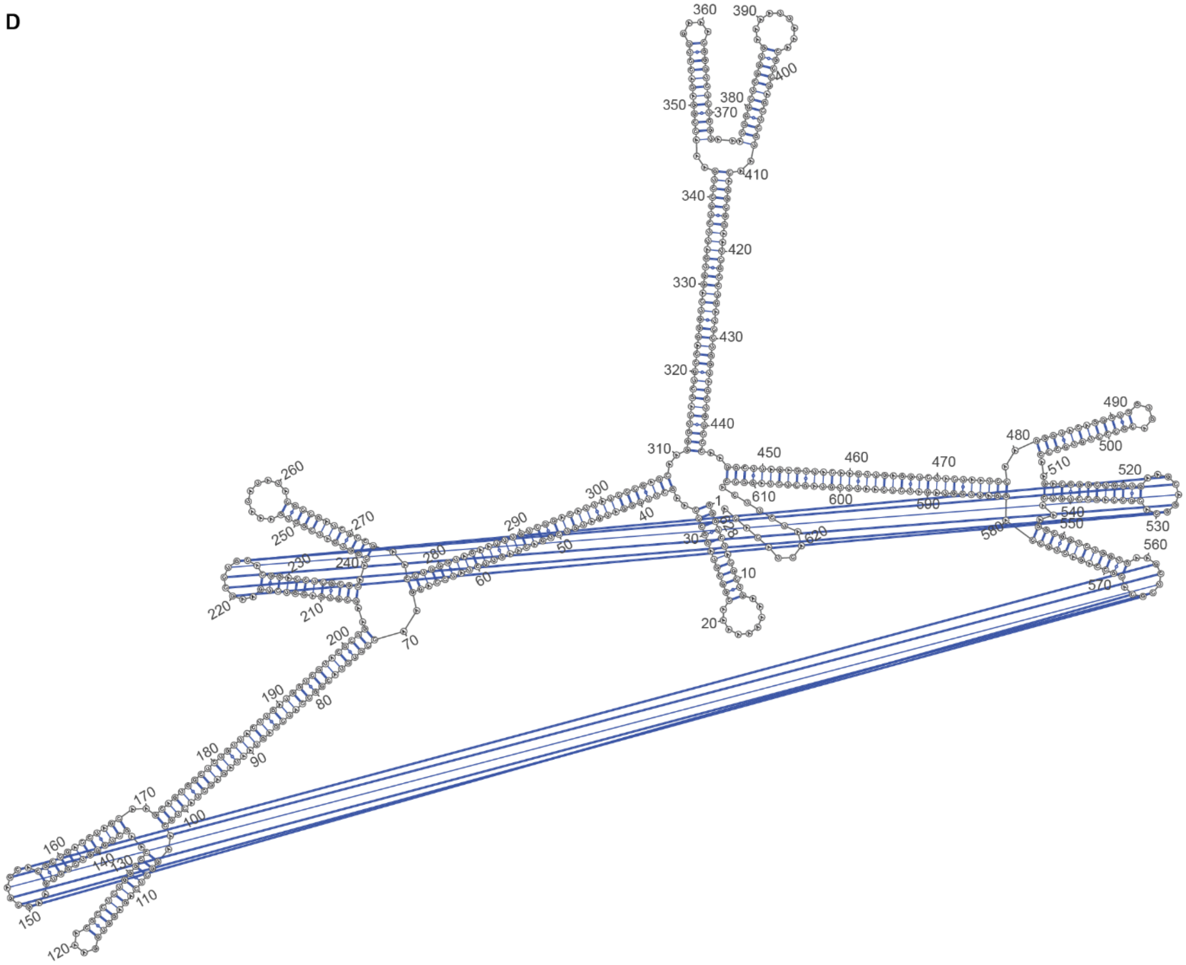
360
390
400
350
370
380
410
340
420
330
430
320
440
310
450
460
470
480
490
500
510
520
530
540
550
560
570
580
590
600
610
620
1
10
20
30
40
50
60
70
80
90
100
110
120
130
140
150
160
170
180
190
200
210
220
230
240
250
260
270
280
290
300

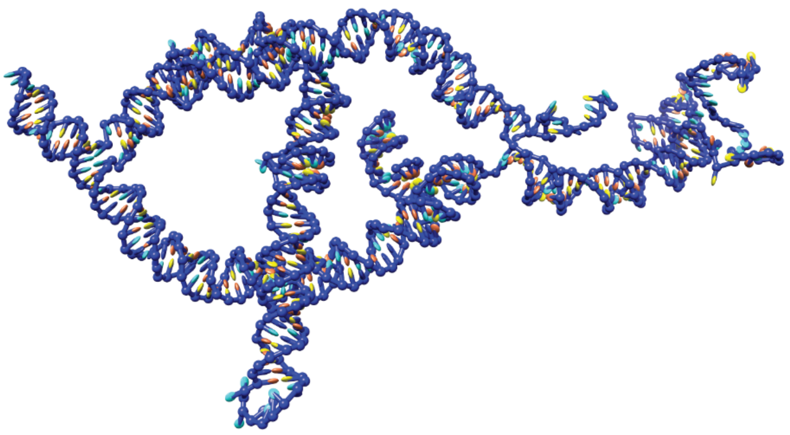

B$_2$

**E**

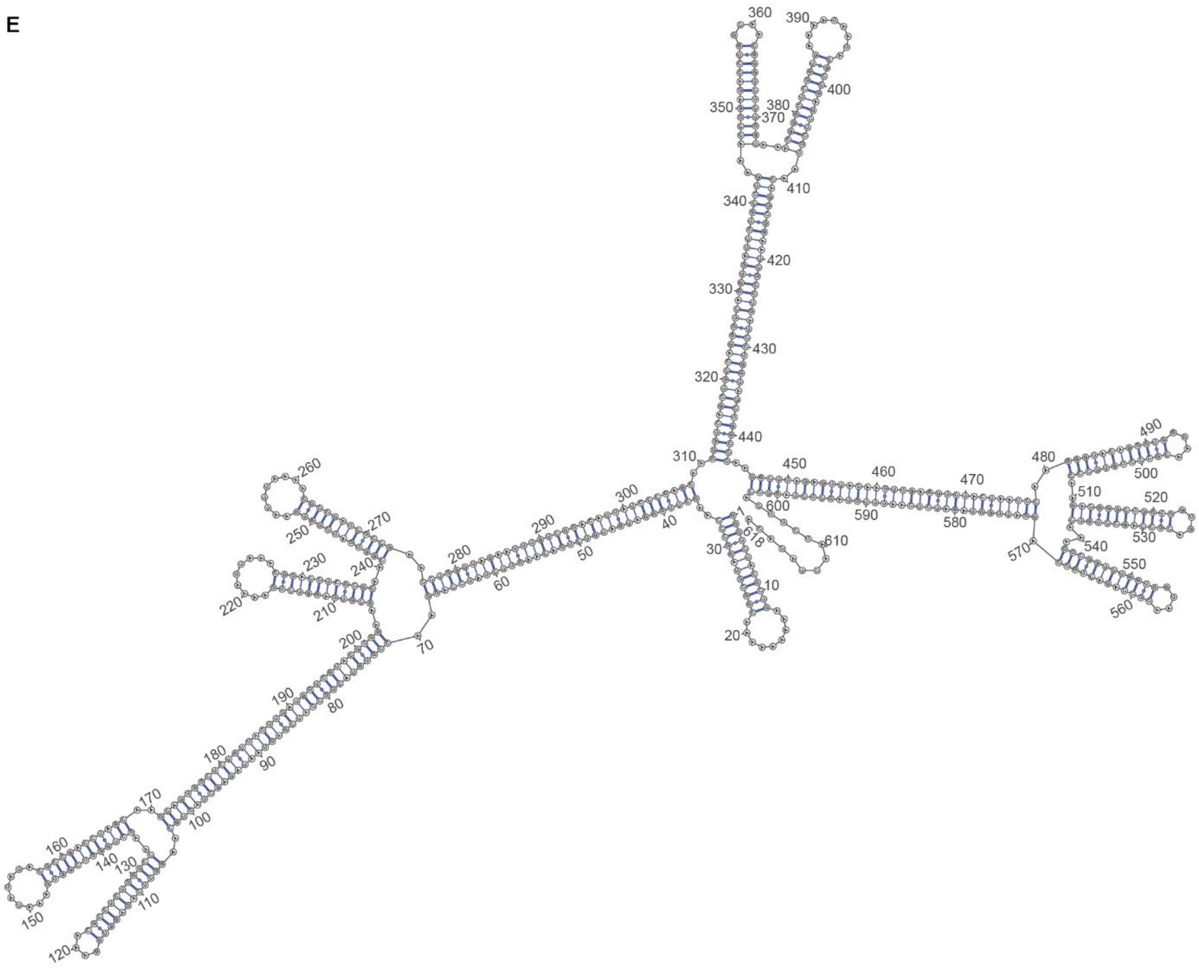
360
390
400
350
380
370
410
340
420
330
430
320
440
310
450
460
470
480
490
500
510
520
530
540
550
560
570
580
590
600
610
618
1
10
20
30
40
50
60
70
80
90
100
110
120
130
140
150
160
170
180
190
200
210
220
230
240
250
260
270
280
290
300

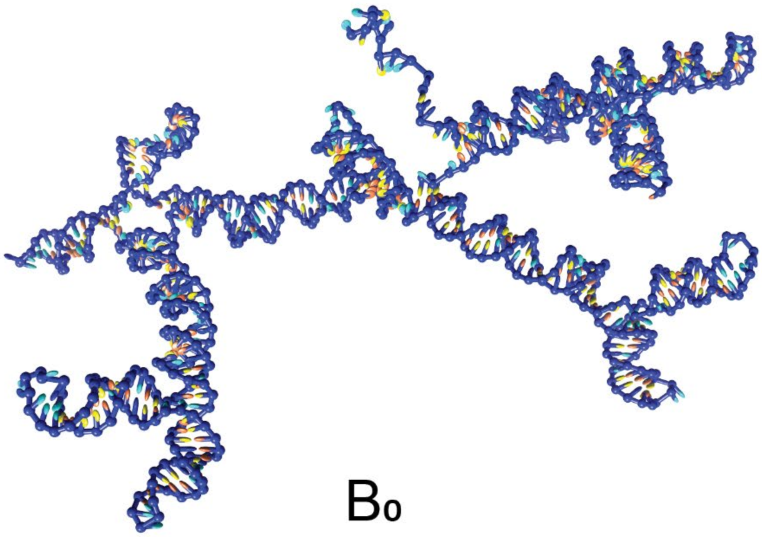
B0

F

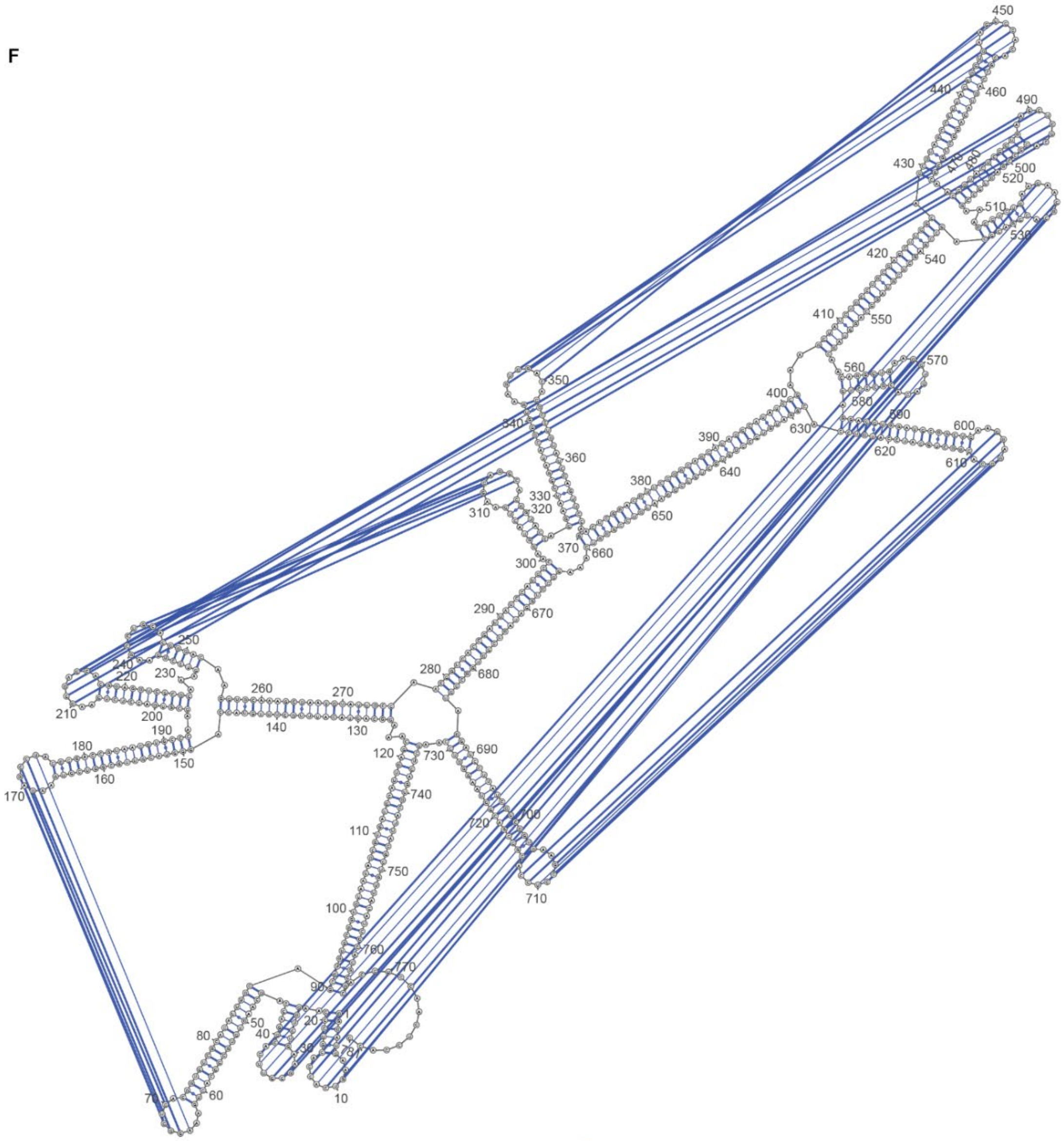
450
440
460
490
430
500
520
510
530
420
540
410
550
570
560
580
590
400
630
600
620
610
350
340
360
390
640
380
650
330
320
310
370
660
300
290
670
250
240
230
220
210
200
190
180
150
170
160
260
270
140
130
280
680
120
730
690
740
110
720
700
710
750
100
760
770
90
80
50
40
20
30
70
60
10

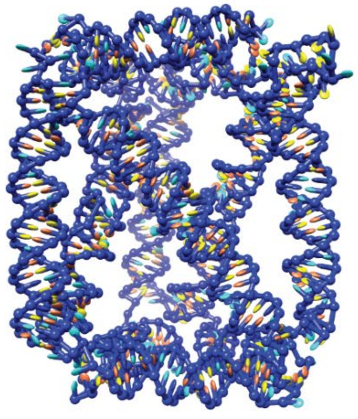

P

**G**

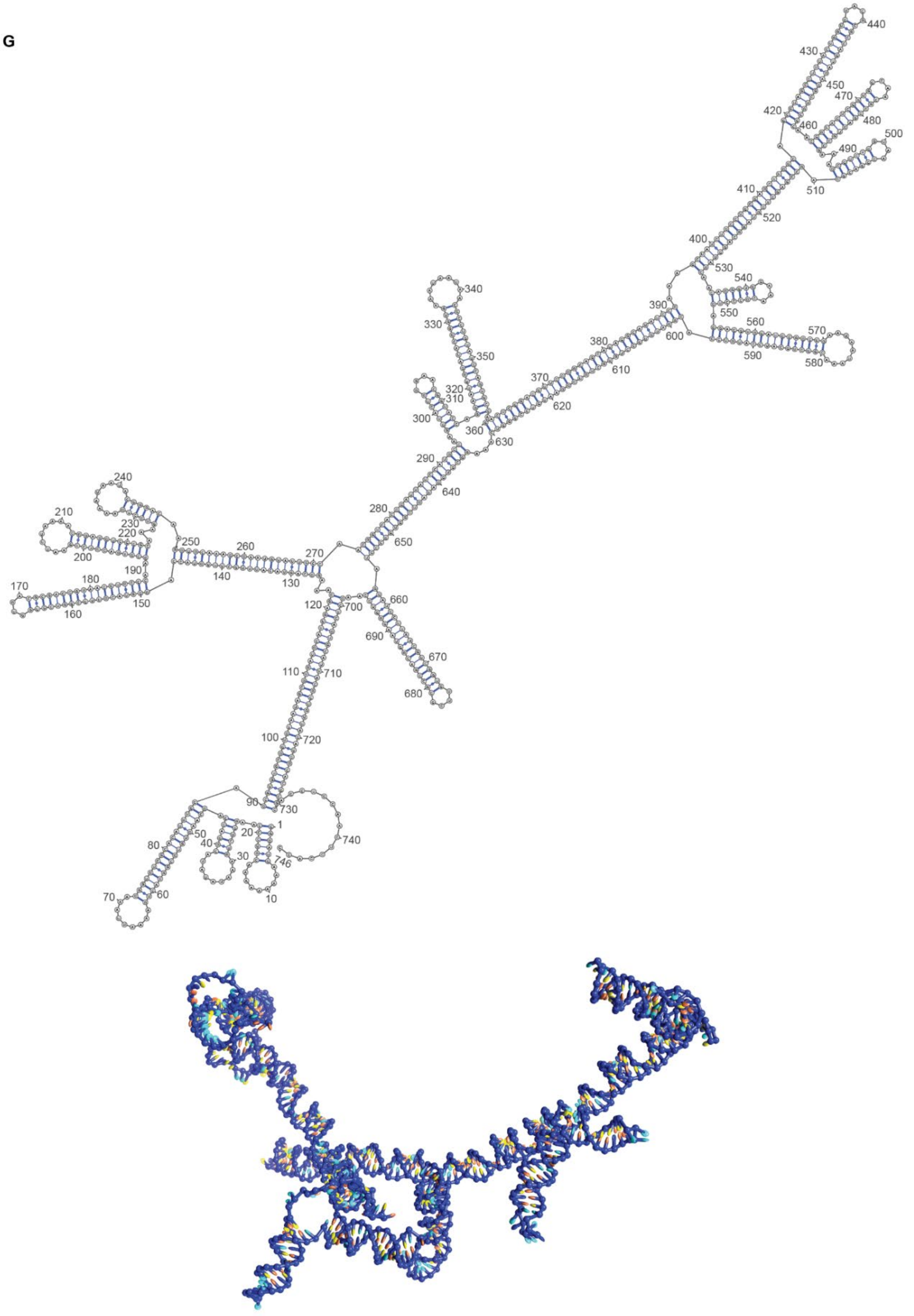


$P_0$

**Fig. S7 Secondary structure diagrams from IPKnot and oxRNA simulation snapshots of RNA structures: (A)** Structure T: Tetrahedron with all 3 kissing loops intact. **(B)** Structure $T_0$: Deficient tetrahedron with all 3 kissing loops disabled. **(C)** Structure B: Bipyramid with all 5 kissing loops intact. **(D)** Structure $B_2$: Deficient bipyramid with 3 out of 5 kissing loops disabled. **(E)** Structure $B_0$: Deficient bipyramid with all 5 kissing loops disabled. **(F)** Structure P: Triangulated prism with all 7 kissing loops intact. **(G)** Structure $P_0$: Deficient prism with all 7 kissing loops disabled.

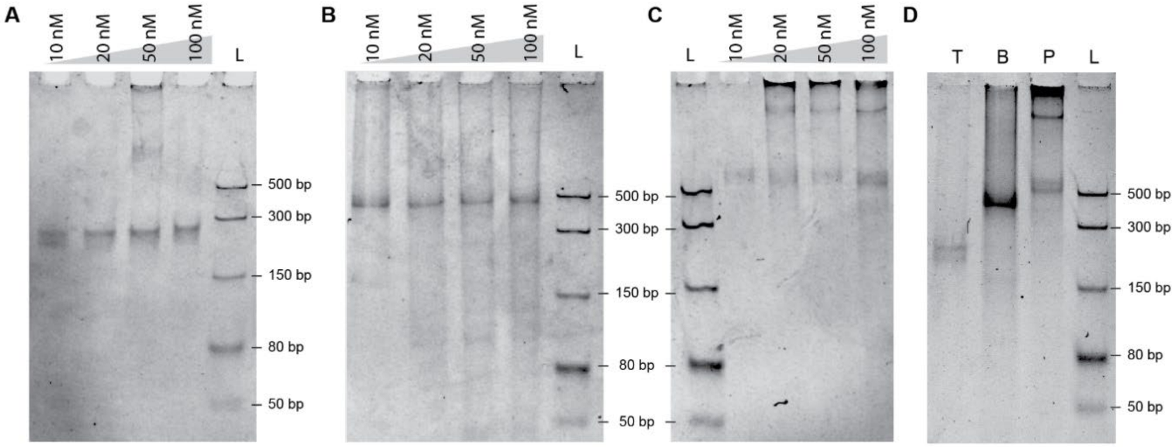


**Fig. S8 Concentration optimization for folding of RNA structures.** Optimization of folding concentration with 2 mM $Mg^{2+}$ analysed using 5% native PAGE of **(A)** Tetrahedron, **(B)** Bipyramid and **(C)** Prism. The samples were diluted to 100 fmoles and added to the wells. L is a dsRNA ladder. Tetrahedron and Bipyramid folded without aggregation, while Prism folded at 10 nM concentration without aggregation. At higher concentrations (≥ 20 nM), severe aggregation seems to occur when folding this structure. **(D)** Comparison of folded RNA nanostructures folded at 50 nM concentration. Tetrahedron and Bipyramid form a single band whereas Prism shows aggregation.

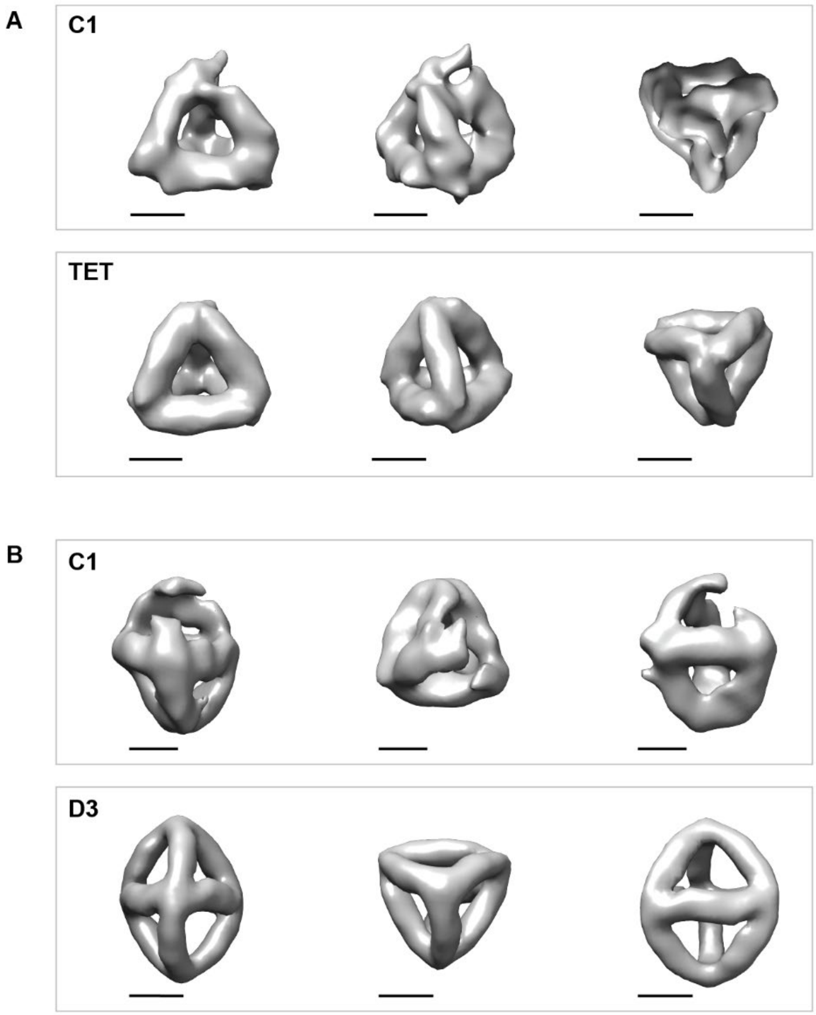


**Fig. S9 Reconstruction of Tetrahedron and Bipyramid with different symmetries. (A)** Refined reconstruction of Tetrahedron with C1 and TET symmetry. **(B)** Refined reconstruction of Bipyramid with C1 and D3 symmetry. Scale bars: 5 nm.

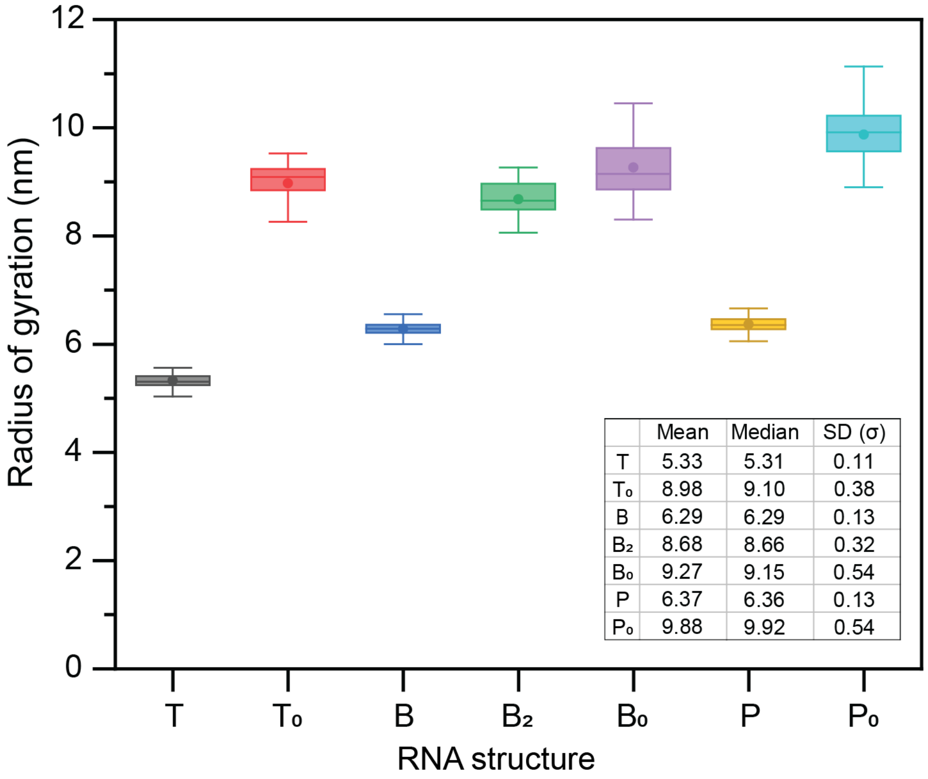


| | Mean | Median | SD (σ) |
|---|---|---|---|
| T | 5.33 | 5.31 | 0.11 |
| $T_o$ | 8.98 | 9.10 | 0.38 |
| B | 6.29 | 6.29 | 0.13 |
| $B_2$ | 8.68 | 8.66 | 0.32 |
| $B_o$ | 9.27 | 9.15 | 0.54 |
| P | 6.37 | 6.36 | 0.13 |
| $P_o$ | 9.88 | 9.92 | 0.54 |

**Fig. S10 Analysis of radius of gyration from oxRNA simulations.** The radius of gyration of RNA structures was evaluated from the results of oxRNA simulation runs using Visual Molecular Dynamics (VMD, https://www.ks.uiuc.edu/Research/vmd/). We used the last 100 frames of the simulations to get the mean (circle), median (horizontal line), inter-quartile range (box) and maximum and minimum values of the radius in nanometres. The last 100 frames were employed to make sure that the simulated structures were relaxed throughout the sample.

**Table S1: P-stick model parameters.** Adapted from the 2014 paper by Geary and Andersen (*21*).

| Parameter | Variable | A-form |
|---|---|---|
| Radius | R | 8.7 Å |
| Rise | D | 2.81 Å |
| Inclination | I | -7.45 Å |
| Axis | A | 139.9 Å |
| Twist | T | 32.73 Å |
| Helicity | H | 11 bp |

**Table S2: The top ten kissing loop pairs from a possible seventeen million combinations of six base kissing loops.** Note: The energies were calculated with the A + AA linkers included around the loops (though not shown in the list).

| Sequence | Binding Energy (kcal/mol) |
|---|---|
| GGGCCG GCCCGG | −13.8840164 |
| GGCCCG GCCGGG | −13.8840164 |
| GGGCCG ACCCGG | −13.8840163 |
| GGCCCA GCCGGG | −13.8840163 |
| GGCCCA ACCGGG | −13.8840163 |
| GGGCCA ACCCGG | −13.8840163 |
| GGGCCA GCCCGG | −13.8840163 |
| GGCCCG ACCGGG | −13.8840163 |
| GCCCCA GCGGGG | −13.8831708 |
| GGGGCG ACCCCG | −13.8831708 |

**Table S3 DNA templates and primer sequences used in the experiments.** T: Tetrahedron with all kissing loops intact. $T_0$: Tetrahedron with no kissing loops. B: Bipyramid with all kissing loops intact. $B_2$: Bipyramid with two kissing loops intact. $B_0$: Bipyramid with no kissing loops. P: Prism with all kissing loops intact. $P_0$: Prism with no kissing loops. FP: Forward primer for all three sequences. RP-T: Reverse primer for Tetrahedron. RP-B: Reverse primer for Bipyramid. RP-P: Reverse primer for Prism.

| | |
|---|---|
| T | GACTAATACGACTCACTATAGGGACGATATGTCCTGCAAGACCGGTCGGT<br>GAAGGAGGCACGCCGACTGGTCAAGCTCGGCTGGTGAAGCGAGCACGCC<br>AGCTGAGCAAGCAGGGCATATTGTCCCGCTCTAAGCCTATGACAAGACTG<br>TACATACGGATTGTGCACCTCTAGCCGCAAGCTGTCCAGGTGAAGCTCGC<br>ACGCCTGGGCAGCAAACGCGGAGGCTGAAGTGGACACGGCCTCTGCGTA<br>AGCGGCTGGAGGTGTACAATCTGTATGTGCAGTCAACGTTCTCGTCTCGG<br>TCAGCTGCACTGGTCGATGAAGCTCTGGGACTGAAGTCCACACGGTCCCG<br>GAGCAACGTCTGGTGCTGAAGCCTCCACGGCACCGGACGAACATCGATC<br>AGTGCGGCTGACTGAGACGGGAACGAAGTCATGGGCTTGGAGCACTCAA<br>CTTCACACC |
| $T_0$ | GACTAATACGACTCACTATAGGGACGATATGTCCTGCAAGACCGGTCGGT<br>GAAGGAGGCACGCCGACTGGTCAAGCTCGGCTGGTGAAAATTAAACGCC<br>AGCTGAGCAAGCAGGGCATATTGTCCCGCTCTAAGCCTATGACAAGACTG<br>TACATACGGATTGTGCACCTCTAGCCGCAAGCTGTCCAGGTGAAGCTCGC<br>ACGCCTGGGCAGCAAACGCGGAGGCTGAAATATATACGGCCTCTGCGTA<br>AGCGGCTGGAGGTGTACAATCTGTATGTGCAGTCAACGTTCTCGTCTCGG<br>TCAGCTGCACTGGTCGATGAAGCTCTGGGACTGAAGTCCACACGGTCCCG<br>GAGCAACGTCTGGTGCTGAAAAAAAAACGGCACCGGACGAACATCGATC<br>AGTGCGGCTGACTGAGACGGGAACGAAGTCATGGGCTTGGAGCACTCAA<br>CTTCACACC |
| B | GACTAATACGACTCACTATAGGGTCTACGCTGAAAGGCTCACGGCGTAGG<br>CCCAAGCTTGGGATGATGTCTGACAAGTTCTATCCAGGAACCGTGTACGG<br>CCATCGAGTAATAGAGCTACTGCAAGGCTCAGAGGTGAAGAGCCTACGC<br>CTCTGGGCCAAGCTGGGTCGGTGAAGCGAGCACGCCGACCTAGCAAGCA<br>GTGGCTCTGTTACTTGATGGTCGTACGCGGAAGCGTGAGGTCTGAAGCCT<br>GCACGGACCTCGCGCAACGGTGATCGGTGAAGTGGACACGCCGATCGCC<br>GAACCTGGGTAGAATTTGTCGGACATTATCCCGAGCAAGGGTCAGCTGCC<br>AGGGTCAGGTGATTCTGCCTGAAACCGGAGACCTGAAGTCCACACGGGTC<br>TCTGGTAAACGGGCTCGGTGAAGGAGGCACGCCGAGCTCGTAACAGGCG<br>GAATCGCCTGATCCTGGTAGCTGGCCCAAGGCTTAGAGTTACAAGTTGAG<br>GTCTACTAATCCAAGGGTACAGGGTGAAGCCTCCACGCCCTGTGCCCAAG<br>ACGGGCTGGTGAAGCAGGCACGCCAGCCTGTCAAAGGCGGTCTGGTGAA<br>GCTCGCACGCCAGACTGCCAGGATTGGTAGATCTCAATTTGTAGCTCTAG<br>GCCACTTCTAACTACACA |
| $B_2$ | GACTAATACGACTCACTATAGGGTCTACGCTGAAAAAAAAACGGCGTAG<br>GCCCAAGCTTGGGATGATGTCTGACAAGTTCTATCCAGGAACCGTGTACG<br>GCCATCGAGTAATAGAGCTACTGCAAGGCTCAGAGGTGGAAACGCCTCT<br>GGGCCAAGCTGGGTCGGTGAAGCGAGCACGCCGACCTAGCAAGCAGTGG<br>CTCTGTTACTTGATGGTCGTACGCGGAAGCGTGAGGTCTGAAGCCTGCAC<br>GGACCTCGCGCAACGGTGATCGGTGAAATATATACGCCGATCGCCGAACC<br>TGGGTAGAATTTGTCGGACATTATCCCGAGCAAGGGTCAGCTGCCAGGGT<br>CAGGTGATTCTGCCTGAAACCGGAGACCTGGAAACGGGTCTCTGGTAAAC<br>GGGCTCGGTGAAAATTAAACGCCGAGCTCGTAACAGGCGGAATCGCCTG |

| | |
|---|---|
| | ATCCTGGTAGCTGGCCCAAGGCTTAGAGTTACAAGTTGAGGTCTACTAAT CCAAGGGTACAGGGTGGTGACGCCCTGTGCCCAAGACGGGCTGGTGAAG CAGGCACGCCAGCCTGTCAAAGGCGGTCTGGTGAAGCTCGCACGCCAGA CTGCCAGGATTGGTAGATCTCAATTTGTAGCTCTAGGCCACTTCTAACTAC ACA |
| $B_0$ | GACTAATACGACTCACTATAGGGTCTACGCTGAAAAAAAAACGGCGTAG GCCCAAGCTTGGGATGATGTCTGACAAGTTCTATCCAGGAACCGTGTACG GCCATCGAGTAATAGAGCTACTGCAAGGCTCAGAGGTGGAAACGCCTCT GGGCCAAGCTGGGTCGGTGAAATATATACGCCGACCTAGCAAGCAGTGG CTCTGTTACTTGATGGTCGTACGCGGAAGCGTGAGGTCTGAAAATTAAAC GGACCTCGCGCAACGGTGATCGGTGAATTTAAAACGCCGATCGCCGAACC TGGGTAGAATTTGTCGGACATTATCCCGAGCAAGGGTCAGCTGCCAGGGT CAGGTGATTCTGCCTGAAACCGGAGACCTGGCAACGGGTCTCTGGTAAAC GGGCTCGGTGAAAATAATACGCCGAGCTCGTAACAGGCGGAATCGCCTG ATCCTGGTAGCTGGCCCAAGGCTTAGAGTTACAAGTTGAGGTCTACTAAT CCAAGGGTACAGGGTGGGAACGCCCTGTGCCCAAGACGGGCTGGTGGTG ACGCCAGCCTGTCAAAGGCGGTCTGGTGGTAACGCCAGACTGCCAGGATT GGTAGATCTCAATTTGTAGCTCTAGGCCACTTCTAACTACACA |
| P | GACTAATACGACTCACTATAGGGATGAAGTCACCACGTCCCAAGGTCTGA AGCGTTCACGGACCAGCAAGTGTGCGGCACTGAAAGGCTCACGGTGTCG CACATTTGCAGCGTGTGCGGTGTAATAGCGTGATGCATTGAGCAAAGGCA GTACATTTGCTTTGTACCAGGGTACGTTGCGGTCGTGAAGAGCCTACGCG ACCGTAACGTGCCCAAGCGTGACCTCTGAAGCGAGCACGGAGGTCGCGC AAAGTCCTGAAGCCTGCACGGGACAAGGTGCAAAGCGAATGTATTGCCA ACGGTCACCGATTATCCATGCCAAGGTACTGAAGCAGGCACGGTACCAG GAATTGGATACGGGTGAAGTGGACACGCCTGTATCCAGTTCCAGACTCGG ACGTTCCGGCAGCATAGGCATATCCGAAAGCTATCTGCCTGCGAGCTTGG CAGCGTAGCCTTTACTGGTGAAGTCCACACGCCAGTAGAGGCTGCGCAAC GGTACTCGGTGAAGCTCGCACGCCGAGTGCCGAAAGCGTCTGAAGAACG CACGGACGCAGCCGAGCTCGTAGGCAGGTAGCAACAGGGTGAAGGTGAC ACGCCCTGAGGACGTGGATGCGGCTGAAGGAGGCACGGCTGCATCCATG TCCACGGATGTGCCTGTGCTGTCGGAATGTCCGGGTCAAAGGCGTGGATA GTCGGTGGCCGTAGGATCCTTATCGTCGGTGAAGCCTCCACGCCGACGGT AAGGGTCCAAGCTCAGTGCATTACGCTGTTACATCGCACGCGCACCTCCA ACTCCACC |
| $P_0$ | GACTAATACGACTCACTATAGGGATGAAAAAAAAACGTCCCAAGGTCTG AAATATATACGGACCAGCAAGTGTGCGGCACTGAAAATTAAACGGTGTC GCACATTTGCAGCGTGTGCGGTGTAATAGCGTGATGCATTGAGCAAAGGC AGTACATTTGCTTTGTACCAGGGTACGTTGCGGTCGTGGTGACGCGACCG TAACGTGCCCAAGCGTGACCTCTGAATTTAAAACGGAGGTCGCGCAAAGT CCTGAAAATAATACGGGACAAGGTGCAAAGCGAATGTATTGCCAACGGT CACCGATTATCCATGCCAAGGTACTGGGAACGGTACCAGGAATTGGATAC GGGTGAAATAATAACGCCTGTATCCAGTTCCAGACTCGGACGTTCCGGCA GCATAGGCATATCCGAAAGCTATCTGCCTGCGAGCTTGGCAGCGTAGCCT TTACTGGTGTACGCGCCAGTAGAGGCTGCGCAACGGTACTCGGTGGCAAC GCCGAGTGCCGAAAGCGTCTGGTAACGGACGCAGCCGAGCTCGTAGGCA GGTAGCAACAGGGTGGAAACGCCCTGAGGACGTGGATGCGGCTGAATAA TAAACGGCTGCATCCATGTCCACGGATGTGCCTGTGCTGTCGGAATGTCC GGGTCAAAGGCGTGGATAGTCGGTGGCCGTAGGATCCTTATCGTCGGTGT |

| | |
|---|---|
| | TCGCGCCGACGGTAAGGGTCCAAGCTCAGTGCATTACGCTGTTACATCGC ACGCGCACCTCCAACTCCACC |
| FP | GACTAATACGACTCACTATAGGGAC |
| RP-T | GGTGTGAAGTTGAGTGCTCC |
| RP-B | TGTGTAGTTAGAAGTGGCCTAG |
| RP-P | GGTGGAGTTGGAGGTGCGC |
